# Proteonano™: A Robust Platform for Large Cohort Deep Plasma Proteomic Studies

**DOI:** 10.1101/2024.08.20.608582

**Authors:** Hao Wu, Yonghao Zhang, Yi Wang, Xiehua Ouyang

## Abstract

Complete profiling of human plasma proteome is an immerse source for disease biomarker discovery. Cutting-edge mass spectrometers, like ThermoFisher’s Orbitrap Astral, have promised unprecedented insights into the exploration of multiple protein biomarkers from human plasma samples. However, large-scale, deep profiling of the human plasma proteome, especially low-abundant proteins (LAPs, <10 ng.mL^-1^), in a robust and fast way remains challenging. This is largely due to the lack of standardized and automated workflows including LAPs enrichment, reduction, and enzymatic digestion procedures. Until now, these complex procedures have not been incorporated into a streamlined workflow to achieve reproducibility, high-throughput, and deep proteome coverage.

Here we report the Proteonano Ultraplex Proteomics (PUP) Platform for large cohort plasma proteomics studies with robustness and fast throughput by standardizing workflow by incorporating the Proteonano platform and high-resolution mass spectrometers, including Orbitrap Exploris™ 480, Orbitrap Astral™, and timsTOF Pro. This pipeline demonstrates excellent stability and reproducibility, with tunable balance between proteome coverage and throughput. We further demonstrate the utility of this platform for potential biomarker discovery in a neurodegenerative disease cohort. This harmonized method enables robust, fast and large-cohort plasma proteomics studies to meet the need to discover new biomarkers.

## Introduction

The development of untargeted bottom-up mass spectrometry-based proteomics has evolved over the past two decades. Proteomic studies have played a pivotal role in identifying disease biomarkers, which are now integral to precision medicine, contributing to disease detection, patient stratification, therapeutic treatment monitoring, and predictive analysis^1–9^. Blood plasma, a rich repository of novel protein biomarkers, presents unique challenges due to the wide dynamic range (>10 orders of magnitude) in protein concentrations. High-abundance plasma proteins often overshadow the detection of medium- and low-abundance proteins during mass spectrometry-based proteomic analyses^10–12^.

Therefore, prior to mass spectrometry, enrichment and fractionation of low-abundance proteins from plasma samples are commonly employed to mitigate the interference caused by high-abundance proteins^10,11,13^. One widely used approach involves the immunodepletion of the most abundant plasma proteins using immunoassay columns, such as the High-Select Top 14 mini columns. Immunodepletion techniques typically enhance protein detection depth, achieving a median 4-fold increase in identified protein groups. However, studies have shown that depleted plasma primarily contains medium- to high-abundance proteins, with only 5–6% identified as low-abundance^12^. Additionally, immunodepletion poses challenges such as non-specific binding leading to unintended loss of low-abundance proteins, complex sample handling, high costs, and batch effects, limiting its scalability for large-cohort plasma proteomics studies^12^.

Another substantial barrier to the widespread adoption of mass spectrometry (MS)-based proteomics in large cohort studies is the reproducibility of distinct mass spectrometry configurations across laboratories. Ruedi et al.^14^ assessed the quantitative reproducibility, dynamic range, and sensitivity of 11 mass spectrometers globally. Their study quantified over 4,000 proteins from HEK293 cells, achieving a median coefficient of variation (CV) of 57.6% before normalization and 22.0% after normalization. For plasma proteomics, sample-related biases also greatly impact data reproducibility and subsequent biomarker discovery. To address this issue, Geyer et al.^15^ developed three quality marker panels to evaluate whether plasma samples were contaminated by coagulation, platelets, or erythrocytes.

The aforementioned challenges have resulted in most MS-based proteomic studies being small and specialized. In contrast, affinity binder technologies, despite their higher cost, have been widely adopted for larger proteomics cohorts^16^. Consequently, there is a pressing need for robust, efficient, and high-throughput MS-based proteomic workflows that are cost-effective and suitable for large-cohort studies.

Recently, the advancement of several cutting-edge technologies has driven substantial progress in MS-based plasma proteomics. These technologies include nanoparticle enrichment, automated sample preparation, and high-throughput mass spectrometry. Firstly, surface-engineered superparamagnetic nanoparticles have emerged as a promising technology for enriching low-abundance plasma proteins^10,11,17–19^. These nanoparticles interact with proteins in biological fluids to form a thin layer known as the protein corona, and extensive studies have shown the critical role of protein corona in nanoparticle pathophysiology^20–27^. Secondly, unlike traditional manual methods such as immunodepletion or chromatographic fractionation, magnetic nanoparticle reagent-based protein capture, combined with peptide identification via mass spectrometry, is well-suited for automation using common liquid handling workstations equipped with magnet modules and additional components such as heater-shaker units. Notably, affinity binder-based protein capture integrated with signal readout through next-generation sequencing (NGS)-based proteomics technologies like Olink PEA has already been widely employed in large cohort plasma proteomic studies, offering highly automated workflows. Lastly, newly commercialized high-resolution mass spectrometers, including ThermoFisher’s Orbitrap Astral, have significantly enhanced detection depth and proteome coverage, achieving a maximum throughput of 180 samples per day per instrument^28^.

Thanks to these advancements, mass spectrometry-based nanoparticle-enriched plasma proteomics has seen remarkable advancements in recent years. Blume et al.^10^ introduced a 5-nanoparticle panel, functionalized with chemical ligands, that successfully identified over 2,000 proteins from 141 plasma samples using an automated sample preparation workflow. However, the reliance on multiple nanoparticles, sample fractionation, and high starting plasma volume imposes limitations on throughput and substantially increases overall costs. To address such challenges, MacCoss and colleagues^11^ developed the Mag-Net enrichment strategy, which leverages strong-anion exchange magnetic microparticles to enrich membrane-bound particles, enabling the profiling of over 4,000 plasma proteins. Nevertheless, the heterogeneity of extracellular vesicles across plasma samples and impact of centrifugation conditions may pose challenges for achieving reproducibility between batches, a topic that receives minimal attention in the manuscript. For instance, Beimers et al.^12^ compared six plasma proteomics methods and reported that the Mag-Net strategy achieved a detected depth of 1,200 protein groups.

While significant progress has been achieved, notable challenges continue to persist. A recent meta-analysis of 2,134 published manuscripts in the proteomics analyses of nanoparticle corona revealed substantial variability in proteome coverage^18^. Gharibi et al.^17^ highlighted this issue by distributing three technical replicates of a protein corona sample to 17 mass spectrometry core facilities across the United States, uncovering notable discrepancies in plasma proteome coverage. They further demonstrated that employing a standardized database search could effectively minimize these discrepancies across distinct sites^19^. Therefore, there is a critical need to establish standardized nanoparticle protein corona workflows encompassing nanoparticle characterization, sample preparation, and automation, along with mass spectrometry instrumentation, data analysis pipelines, and achieving an optimal balance between proteome coverage and throughput.

In this study, we introduced the Proteonano Ultraplex Proteomics (PUP) Platform, a comprehensive system integrating proprietary monodispersed magnetic nanoparticle-based reagents for low-abundance protein enrichment, an automated sample preparation workstation for parallel sample preparation (Nanomation G1), and high-resolution mass spectrometry instruments for measurement, as well as built-in quality control procedures to ensure consistent and reliable data monitoring throughout the workflow. The PUP platform was rigorously evaluated for key features, including proteome coverage, the balance between protein depth and throughput, compatibility with various mass spectrometry configurations, and batch-to-batch quantitative reproducibility. Notably, it demonstrated the capacity to profile up to 6,000 proteins from pooled human plasma samples, with an intra-batch quantitative median CV of 15%. Its versatility was further tested across ten LC-MS/MS systems, showcasing its ability to maintain a balance between throughput and proteome coverage. Finally, the platform’s feasibility was validated in a cohort of participants with cognitive impairment, demonstrating its potential for novel biomarker discovery in plasma proteomics studies. To our knowledge, this study presents a fully integrated, robust, and standardized mass spectrometry-based deep plasma proteomics platform. We believe this innovation holds great promise for fostering the widespread adoption of nanoparticle-enriched MS-based platforms, thereby accelerating advancements in plasma biomarker discovery on a broader scale.

## Results

### Overview of the Proteonano workflow

The Proteonano platform is intended to be a standardized, high-throughput solution for large-cohort plasma studies, combining robustness with efficiency. It integrates proprietary plasma proteome enrichment reagents, the Nanomation G1 automated workstation, and high-resolution LC-MS/MS systems (Fig. 1). The reagents feature hierarchically structured peptide-conjugated nanoparticles (PCNPs) to selectively enrich low-abundance proteins while depleting medium- and high-abundance proteins during sample preparation. Details of peptide properties are included in the Supplementary Table 1. The Nanomation G1 workstation includes a heater-shaker module for nanoparticle-plasma incubation, a magnetic module for protein corona separation, and an 8-channel liquid handler for parallel pipetting. The top view of the workstation layout can be seen in the Supplementary Information Fig. S1). For peptide identification, the platform incorporates three high-resolution mass spectrometers—Orbitrap 480, Astral instruments (ThermoFisher Scientific), and timsTOF Pro 2 (Bruker Corporation)—ensuring compatibility with widely established proteomics infrastructure (Fig. 1)

**Figure 1.**
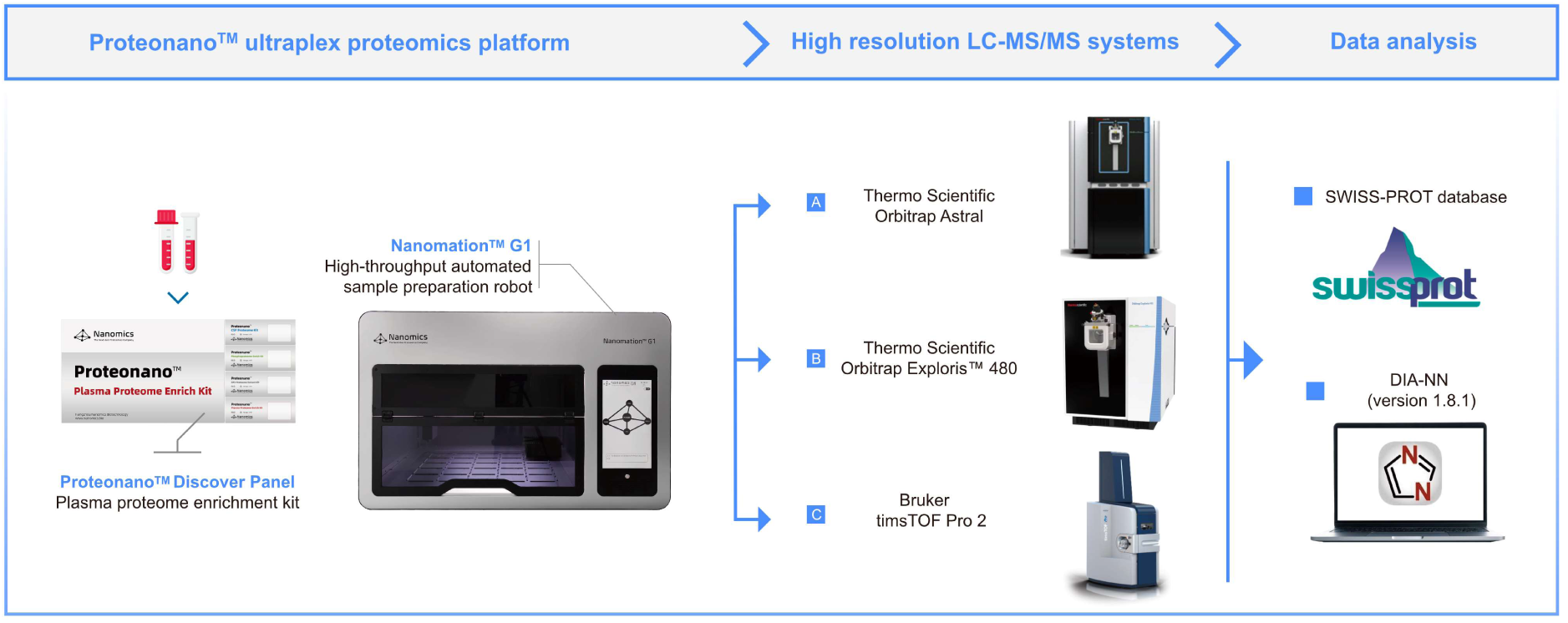
An overall view of the Proteonano Ultraplex Proteomics platform.

The generation of mass spectrometry-based proteomic data is often influenced by various technical and nontechnical factors. To address these challenges, we introduce the Quality Control System (QCS) for the Proteonano workflow—a framework designed to evaluate and ensure data quality and reproducibility. As illustrated in Fig. 2, the QCS encompasses three key control steps: the **Incubation Control** characterizes the proteome enrichment process via nanoparticle-plasma incubation; the **Detection Control** assesses and monitors systematic variations from LC-MS/MS instruments using premade peptide samples; and the **Data Control** includes algorithms for missing data imputation and normalization are employed to detect and correct batch-to-batch variations effectively.

**Fig. 2:**
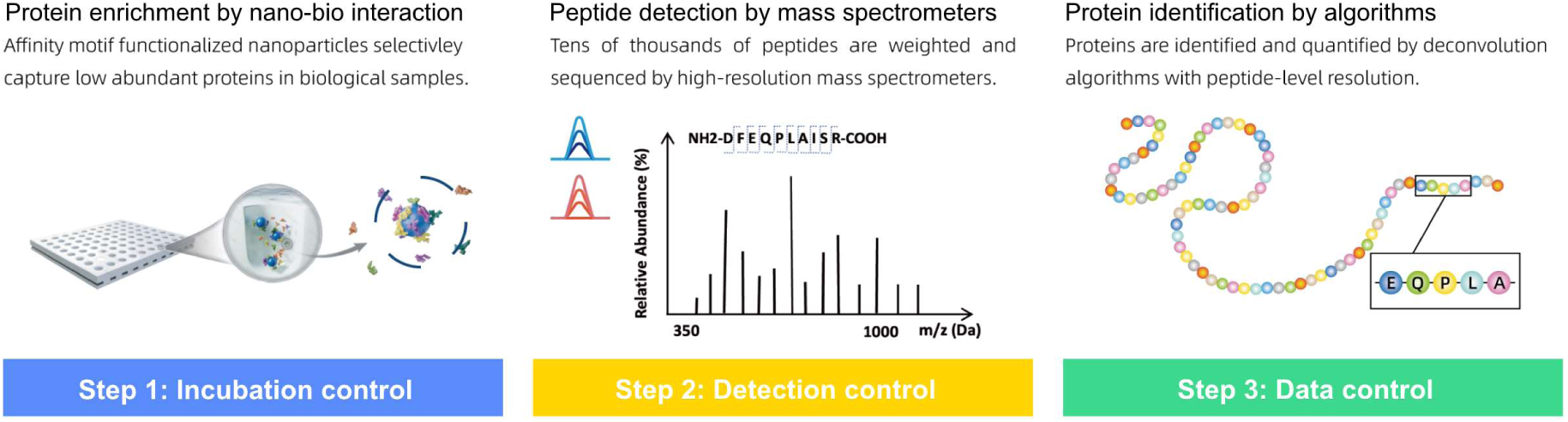
The Quality Control System of the Proteonano Platform for batch-to-batch data monitoring.

In a 96-well plate experiment, six QC samples are used to monitor workflow performance. QC 1 consists of a lyophilized peptide mix derived from pooled plasma, processed with nanoparticle enrichment followed by conventional proteomic preparation steps. It is included in triplicate per plate to assess LC-MS/MS reliability. QC 2 uses neat plasma, untreated by nanoparticles, that undergoes standard sample preparation to evaluate the quality of conventional procedures. QC 3 employs pooled plasma processed nanoparticle enrichment and standard preparation steps to monitor the performance of the complete pipeline. By comparing results from replicates of QC1, QC2 and QC3, the protein enrichment process’s performance can be evaluated. The recommended positions of the six QC samples on the 96-well plate can be seen in the Supplementary Fig. S2.

## Preparation and Characterization of Nanoparticles

The Proteonano nanoparticle is designed to enrich low-abundance proteins from biological samples through nano-bio interactions. As illustrated in Fig. 3a, photographs depict the dispersion of peptide-conjugated nanoparticles in water both before and after peptide conjugation. To fully characterize the multi-layered structures of the PCNPs, we conducted comprehensive measurements on key features, including size, size distribution, surface charge, elemental distribution, and surface ligand density. These analyses were achieved using a combination of advanced techniques including dynamic light scattering (DLS, Fig. 3b, d-g), transmission electron microscopy (TEM, Fig. 3h-n),, zeta potential measurements (Fig. 3c), and high-angle annular dark-field imaging (HAADF-STEM, Fig. 3o-t), and fluorescein isothiocyanate analysis (FTIC, Fig. 3u). The experimental details can be found in the Methods section.

**Figure 3.**
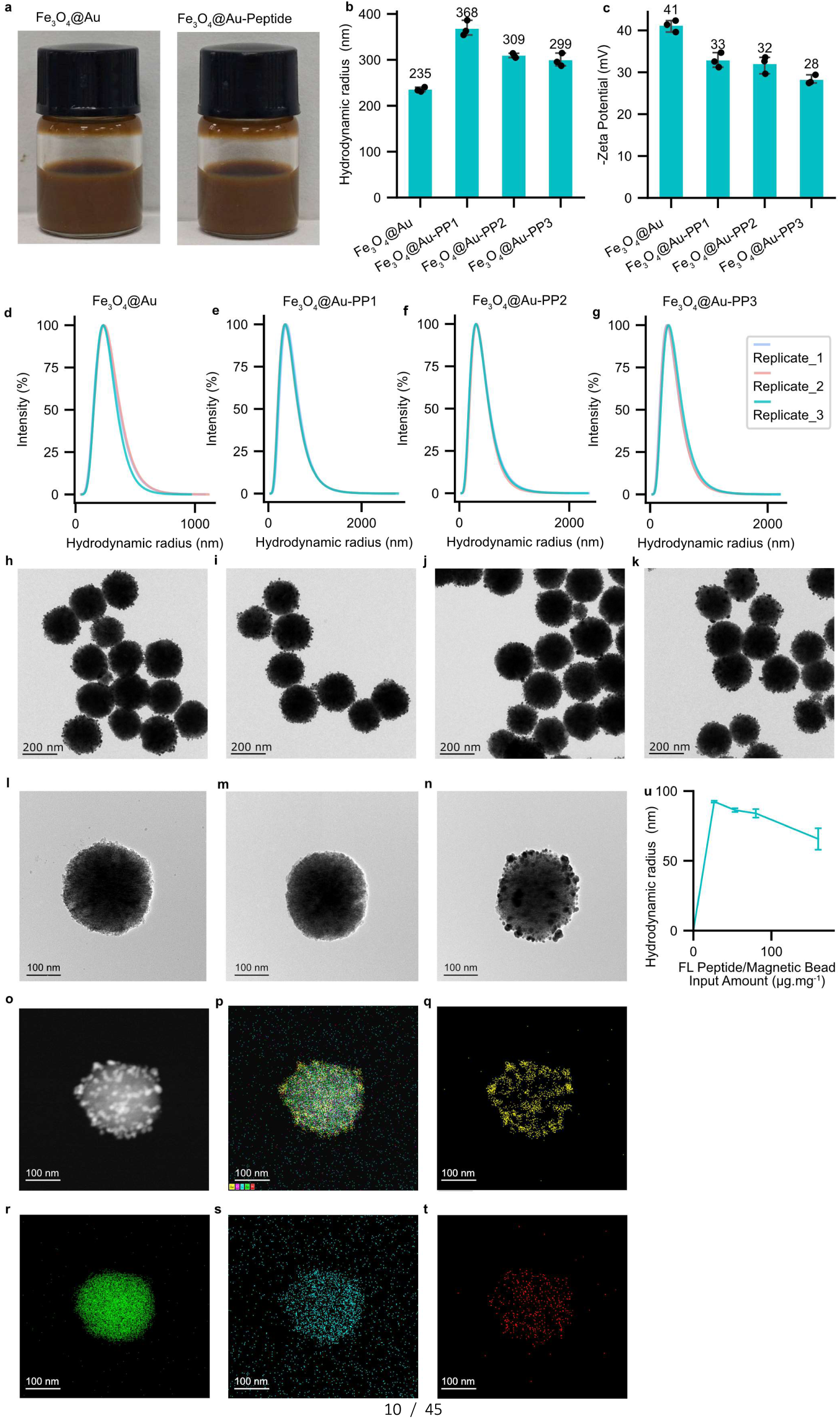
Characterization of peptide-conjugated nanoparticles (PCNPs). **a** Photos of the PCNPs reagent (right) and its precursor (left). **b-k** The hydrodynamic radius and zeta potential (**b**), size distribution (**d-g**), and TEM images (**h-k**) of the Fe3O4@PDA@Au, Fe₃O₄@PDA@Au-PP1, Fe₃O₄@PDA@Au-PP2, and Fe₃O₄@PDA@Au-PP3. **i-n** Representative TEM images of nanoparticles Fe₃O₄, Fe₃O₄@PDA, and Fe₃O₄@PDA@Au. **o Representative** HAADF-STEM images Fe₃O₄@PDA@Au. **p-t** EDS elemental mapping of the dispersion of all elements combined (**p**) and individual element Au (**q**), Fe (**r**), C (**s**) and N (**t**). **u** FITC characterization showing the peptide conjugation density on the surface of nanoparticles.

First, TEM, DLS and zeta potential measurements were utilized to characterize three peptide-conjugated nanoparticles, namely Fe₃O₄@PDA@Au-PP1, Fe₃O₄@PDA@Au-PP2, and Fe₃O₄@PDA@Au-PP3, as well as their precursor (Fe₃O₄@PDA@Au). The Fe₃O₄ nanoparticles exhibit a relatively coarse surface and a regular spherical morphology with an average size of 200 nm (Fig. 3b), which is in line with the hydrodynamic radius and size distribution shown in Fig. 3b, d. A thin layer of polydopamine (PDA) was coated on the surface of Fe₃O₄ nanoparticles to stabilize it. Upon peptide modification, the particle size increased to approximately 300 nm (Fig. 3b, e-g, i-k). The surface properties of the PCNPs were further assessed using zeta-potential measurement. Initially, the nanoparticles displayed a negative zeta-potential of -41.09 ± 1.38 mV, which shifted to around -30 mV following peptide modification, indicating changes in surface charge due to peptide conjugation (Fig. 3c).

Second, HAADF-STEM imaging (Fig. 3l–o) and EDS elemental mapping were performed to investigate the nanoscale structures and elemental distributions of the PCNPs. HAADF-STEM images (Fig. 3l, m, n) show the progression from Fe₃O₄ nanoparticles (l) to Fe₃O₄@PDA nanoparticles (m) to Fe₃O₄@PDA@Au nanoparticles (n). A noticeable surface smoothing was observed after coating Fe₃O₄ with a thin PDA layer. The subsequent attachment of Au onto Fe₃O₄@PDA was evident as discrete nanospheres, rather than a continuous Au layer. EDS elemental maps (Fig. 3q–t) reveal that Au (Fig. 3q) and N (Fig. 3t) are predominantly located on the nanoparticle’s outer surface, while Fe (Fig. 3r) and C (Fig. 3s) are distributed throughout. The combined elemental map (Fig. 3p) and the analysis effectively validate the structure and composition of the synthesized nanoparticles.

Third, the density of peptide ligands on the outer layer was quantified using FITC. As shown in Fig. 2u, at an initial FITC-polypeptide concentration of 24.70 µg.mg^-1^, the input level of fluorescent polypeptide was relatively low, yet the binding efficiency to magnetic beads was high, achieving a conjugation rate of 93%. As the FITC-polypeptide concentration increased, the effective binding efficiency gradually declined. At 53 µg.mg^-1^—consistent with the ligand density on the NP surface (PP1, PP2, PP3)—the conjugation rate decreased to 86%, yielding an estimated ligand density of approximately 46 µg.mg^-1^.

In conclusion, PCNPs can be visualized as a composition of three functional layers. The core layer consists of superparamagnetic Fe₃O₄ nanoparticles with a uniform diameter of 200 nm and remarkable homogeneity. The middle layer features a thin coating of Au particles, deposited as discrete nanospheres. The outer layer comprises chemically modified peptide ligands with a density of approximately 46.0 µg.mg^-1^. These features collectively endow each nanoparticle with excellent size uniformity and a high density of binding sites, enabling the selective enrichment of low-abundance proteins within an extremely small volume. Simultaneously, high-abundance proteins are depleted during sample preparation, facilitating the detection of peptides from rare proteins in subsequent MS measurements.

## Assessment of the plasma proteome coverage

To assess the proteomic detection depth of plasma samples processed with the Proteonano platform, pooled healthy donor plasma samples were analyzed using both the conventional neat plasma pipeline and the Proteonano workflow, followed by LC-MS/MS on the Orbitrap Astral instrument. As shown in Fig. 4a, the Proteonano workflow significantly improved plasma proteome coverage, identifying nearly 4000 protein groups compared to 900 with the conventional pipeline—a ∼330% increase. Among these, ∼2800 protein groups overlapped with proteins cataloged in the Human Plasma Proteome Project (HPPP), whereas only ∼800 overlapped with the neat plasma pipeline results (Fig. 4b). Additionally, the Proteonano workflow detected 123 FDA-cleared circulating plasma biomarkers spanning eight orders of magnitude from a single pooled plasma sample, markedly outperforming the neat plasma pipeline (Fig. 4c).

**Figure 4.**
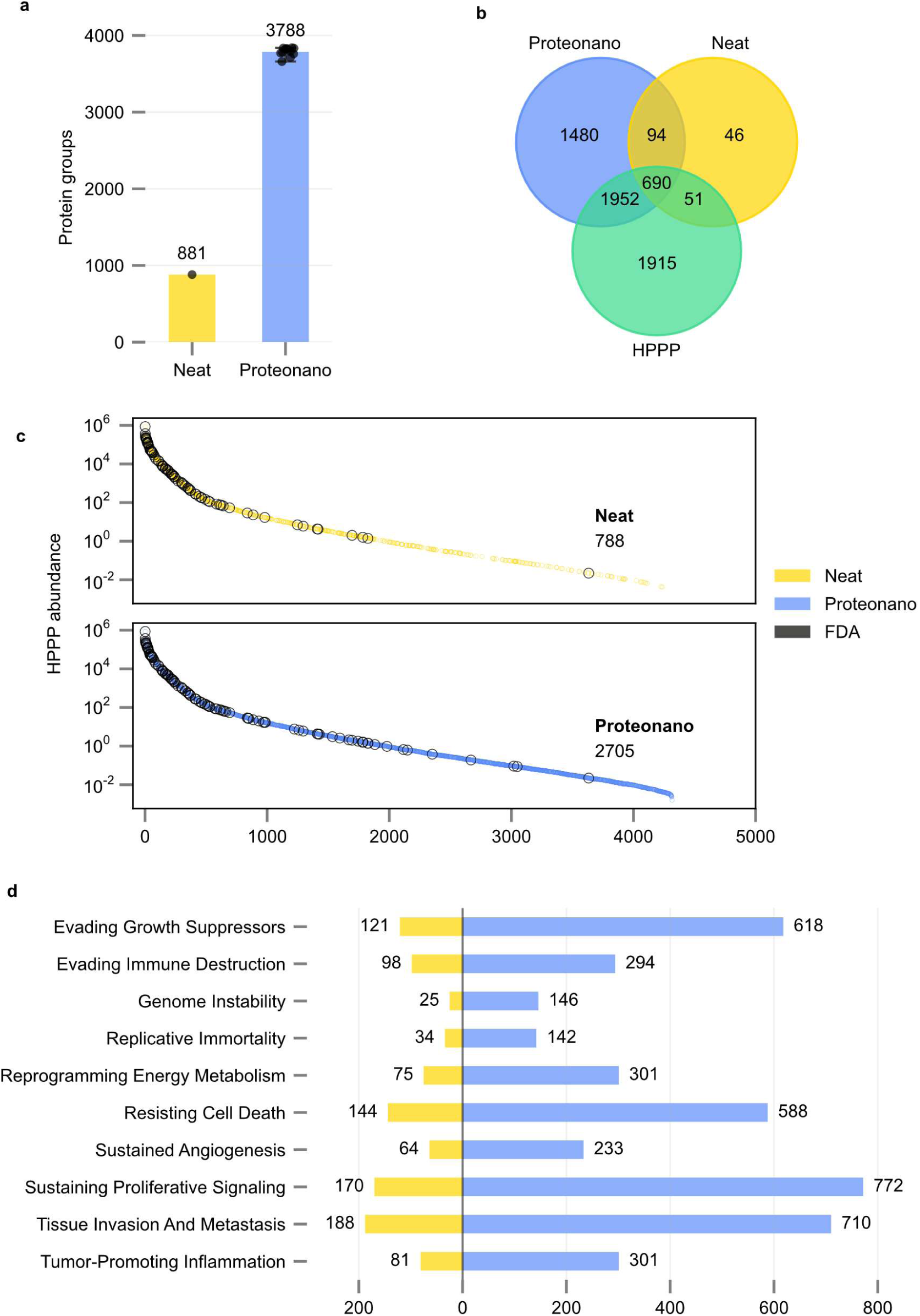
Proteomic coverage of the Proteonano workflow in comparison to neat digestion workflow. **a** Protein groups identified from a pooled healthy donor plasma sample processed using the Proteonano workflow and the neat plasma pipeline. DIA LC-MS/MS measurement was conducted on an Orbitrap Astral instrument (1% protein and peptide FDR; details in the Methods section). **b** Venn diagram comparing plasma proteomic coverage, highlighting overlaps between protein groups identified by the two workflows and those cataloged in the HPPP. **c** Intensity distribution of proteins groups detected matching the HPPP database. Intensities for proteins from neat plasma are shown in the upper panel. Intensities for the Proteonano are shown in the bottom panel. Black circles indicated FDA-approved protein biomarkers. **d** A comparison of the Proteonano workflow and the neat workflow regarding the coverage of secreted proteins associated with 10 hallmarks of cancer progression.

Given the plasma proteome’s composition of secreted proteins from various tissues—both healthy and diseased—it serves as an ideal matrix for disease biomarker discovery and pathogenesis research^29^. We further investigated the secreted proteins linked to the hallmarks of cancer, ten defining traits of cancerous cells: evading growth suppression, immune destruction, genome instability, replicative immortality, reprogrammed energy metabolism, resisting cell death, sustained angiogenesis, proliferative signaling, tissue invasion and metastasis, and tumor-promoting inflammation^30,31^. As shown in Fig. 4d, for each hallmark, the Proteonano workflow identified 3–5 times more secreted proteins compared to the unenriched process. The broader coverage of the plasma secretome associated with cancer progression may position the Proteonano platform as a promising proteomic strategy for early diagnosis, subtyping, and prognosis of various cancers.

## Balance between proteome coverage and throughput on ten LC-MS/MS systems

Besides proteome coverage, throughput (typically defined as samples per day, SPD) is a crucial consideration when adopting proteomics platforms, as it directly affects the speed and scalability of data generation. High-throughput platforms are essential for large-scale or time-sensitive studies, such as biomarker discovery or population analyses. However, there is often a trade-off between throughput and proteome coverage; some platforms prioritize speed, while others focus on achieving comprehensive proteomic analysis at a slower pace. Selecting the right LC-MS/MS system depends on the specific goals, budget, and requirements of the study.

To demonstrate the versatility of the Proteonano workflow, we assessed its coverage-throughput balance across 10 distinct LC-MS/MS systems. These systems featured various combinations of liquid chromatography columns and high-resolution mass spectrometry instruments. Detailed specifications are provided in Supplementary Information Table S2.

We first evaluated the plasma proteome coverage of the Proteonano workflow using the Orbitrap Astral instrument paired with three widely used LC columns (ES906, ES75550, μPac 110 cm), varying throughput from 180SPD to 7SPD. As illustrated in Fig. 5a, even at the shortest gradient (180SPD), the workflow identified 3168±9 protein IDs (AVG±SE, n=3) from a single pooled healthy plasma sample. Gradual increases in gradient time enhanced proteome coverage, yielding 4945±5 protein IDs (AVG±SE, n=3) at 24SPD and 6410±4 protein IDs (AVG±SE, n=3) at 7SPD. Furthermore, with MS/MS parameters kept constant, the proteomic detection depth was evaluated using the ES75500 and μPac 110 cm long LC columns. At comparable throughputs (14SPD), both columns exhibited similar numbers of detected protein groups, indicating the stability of the Proteonano workflow.

**Figure 5.**
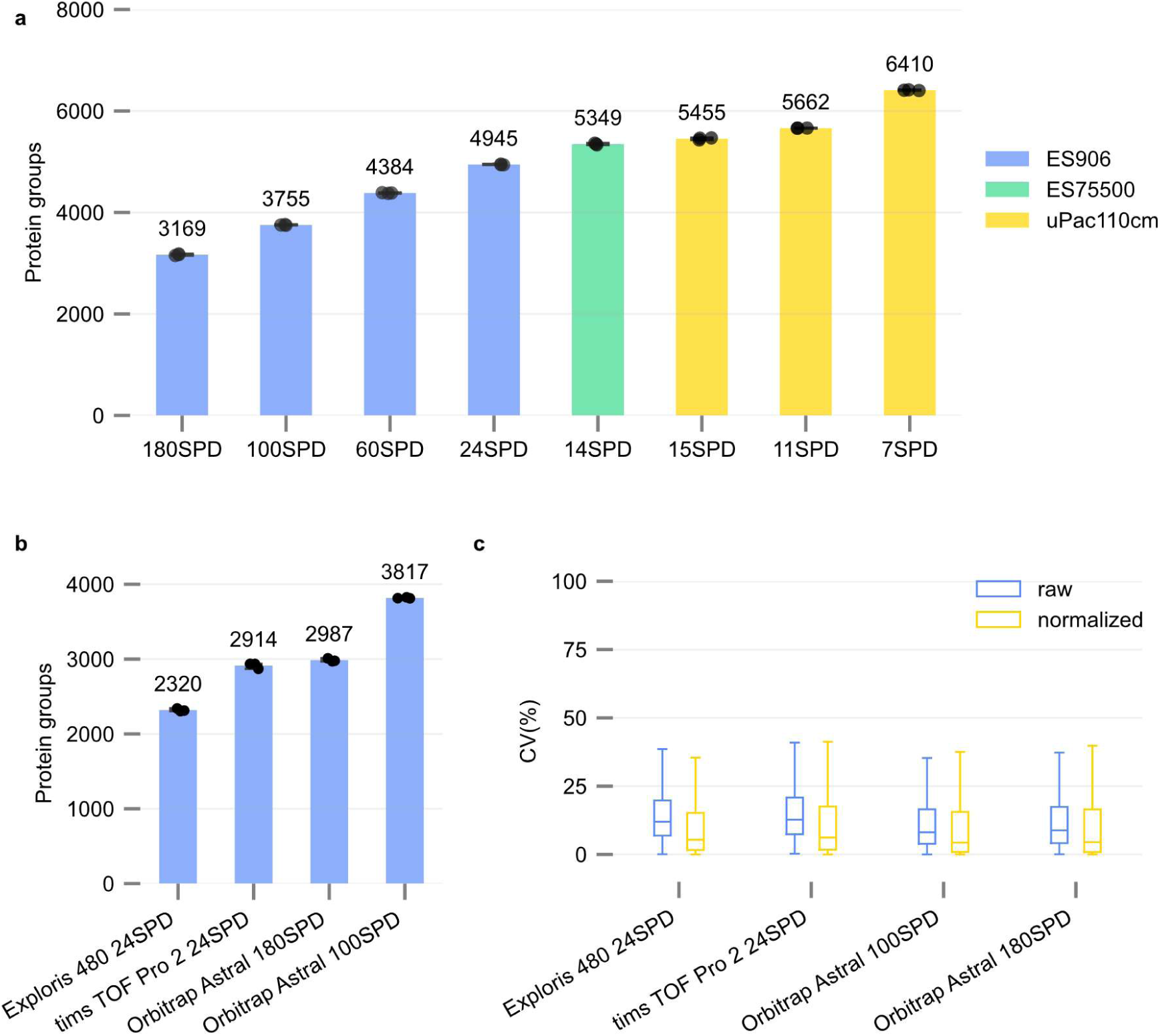
Tunability of proteome coverage and throughput across 10 distinct LC-MS/MS systems. **a** Number of proteins groups identified using different LC columns coupled with the Orbitrap Astral Instrument (DIA mode, 1% protein and peptide FDR). Bar height indicates the mean protein groups detected of three replicates of a pooled plasma sample. Individual dots represent protein groups detected in each replicate. **b** Venn diagram of protein groups of identified in A. **b** Proteome coverage of different mass spectrometry coupled with the same ES906 liquid chromatography column. **c** Distribution of CV values of protein intensity across different LC-MS/MS systems described in **b**.

We further evaluated the impact of the Proteonano workflow on proteomic detection depth and throughput using three mass spectrometers—Orbitrap Exploris 480, Orbitrap Astral, and timsTOF Pro 2— all coupled with ES906 liquid chromatography. Technical assessment was conducted using three replicates of a pooled healthy plasma sample. As shown in Fig. 5b, at 24SPD, Orbitrap Exploris 480 identified 2,320 ± 11 protein IDs (AVG ± SE, n=3), while timsTOF Pro 2 detected 2,914 ± 22 protein IDs (AVG ± SE, n=3). Orbitrap Astral exhibited the best performance, identifying 2,987 ± 12 protein groups (AVG ± SE, n=3) at 180SPD and 3,817 ± 4 protein groups (AVG ± SE, n=3) at 100SPD. All mass spectrometers showed low and similar inter-sample quantitative CVs, with Orbitrap Astral achieving slightly lower CVs (8.10% at 100SPD and 8.80% at 180SPD) compared to timsTOF Pro 2 (12.71% at 24SPD) and Orbitrap Exploris 480 (11.97% at 24SPD) (See Fig. 5c).

These results demonstrate the excellent stability of the Proteonano workflow across various LC-MS/MS systems. Additionally, its tunable and broad depth-throughput bandwidth, ranging from over 3000 protein IDs at 180SPD and up to 6400 at 7SPD, makes it a desirable MS-based platform for deep plasma profiling across different instrument configurations and study requirements.

### Technical Reproducibility of Proteonano Workflow

Affinity-based, non-MS multiplexed proteomics platforms have established comprehensive quality control measures as a fundamental approach to ensure data reproducibility. For instance, Olink implements a rigorous quality control system built on its Proximity Extension Assay (PEA) technology, incorporating validated biomarker panels and automated processes to reduce manual handling while maintaining data integrity. These measures include protein expression normalization and stringent panel performance validation^32^. Similarly, SomaLogic adopts a Total Quality Management approach focused on safety, efficacy, and reproducibility. Their quality control framework integrates normalization techniques, calibration, and adaptive signal adjustments to ensure consistent and accurate proteomic data^33^.

We first evaluated whether variations between PCNP batches influenced proteomic analysis, as reproducibility of the enrichment reagent is vital for quality control. Supplementary Fig. 4 shows high pairwise Pearson correlation coefficients between two batches, confirming the strong reproducibility and consistency of PCNPs in enrichment-based plasma proteomics workflows.

We next assessed the baseline reliability of the LC-MS/MS system (Vanquish Neo liquid chromatography loaded with ES906 column, coupled with Orbitrap Astral) using 12 equal aliquots of the QC1 plasma sample from six batches, which were processed in parallel with the Proteonano workflow. The resulting freeze-dried peptides were dissolved, pooled, and consecutively injected for LC-MS/MS measurements (detailed in the Methods section). As shown in Fig. 6a and 6b, 3641±15.9 protein groups (AVG±SE, n=12) were identified, with an inter-plate coefficient of variation (CV) of 7.69% before and 3.38% after quantile normalization, based on the number of identified protein groups. We further compared the technical CVs of the low abundance proteins versus the medium- to high-abundance proteins. As shown in the boxplot in Fig. 6c, low-abundance proteins displayed higher median CVs (6.65%, <10 ng.mL^-1^) compared to medium-(4.32%, 10–100 ng.mL^-1^) and high-abundance proteins (3.86%, ≥100 ng.mL^-1^). It is anticipated that the higher abundance proteins would be more quantitatively reproducible, as there is a lower noise-to-signal ratio for more reliable quantification.

**Figure 6.**
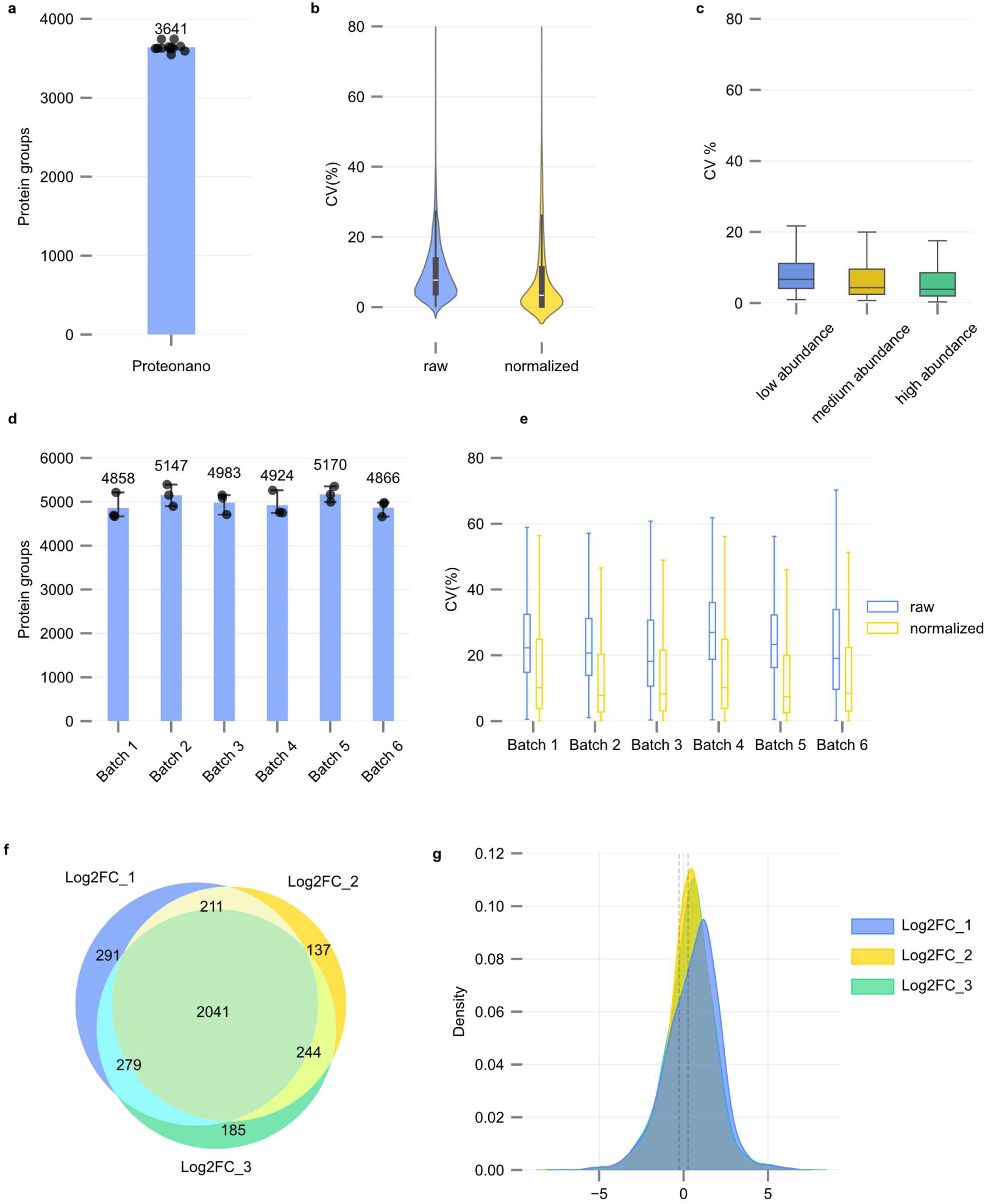
Investigation on the batch-to-batch reproducibility of the Proteonano workflow. **a** Baseline LC-MS/MS system reliability—number of identified protein groups (AVG±SE, n=12). **b** Inter-plate CVs before and after quantile normalization. **c** Technical variability comparison—boxplot illustrating median CVs of low-, medium-, and high-abundance proteins. **d** Inter-batch protein identification consistency—average number of protein groups across six independent batches. **e** Median CVs across batches and assessment of sample preparation variability. **g** Differentially expressed proteins (DEPs)-pairwise comparisons of Log2FC groups across two independent projects. **f** Evaluation of reproducibility using Jaccard Similarity Index.

Second, we evaluated the inter-batch reproducibility of the complete Proteonano workflow across six independent batches with each batch including three replicates of QC3 sample. As the results shown in Fib. 6d and 6e, the average number of protein groups identified in each batch kept relatively consistent, ranging from the lowest of 4858 to the highest of 5170. We next compared the median CV values of the 6 batches. The median CV of the raw unadjusted intensities were in the range of 18.2% to 26.9%. After quantile normalization, the median CVs dropped to 7.4% to 10.2%. Compared to the CVs in Fig. 6c, we noticed a clear increase, which is probably due to the additional variations introduced by the sample preparation steps prior to mass spectrometry measurements. It is reasonable to infer that the coefficient of variation for the complete Proteonano workflow (Fig. 6e) represents the combined variability stemming from the sample preparation process and the LC-MS/MS measurement (Fig. 6c). Based on this rationale, the average CV attributed to the sample preparation procedure was determined to be 16.5%.

Third, the reproducibility of differentially expressed proteins (DEPs) across two independent projects was assessed. Project 1 comprised six batches, each containing three replicates of the QC3 sample (P1QC3), while Project 2 included three batches, each with three replicates of a different QC3 sample (P2QC3). Nine P1QC3 samples were randomly divided into three groups, with the same grouping applied to the nine P2QC3 samples. Pairwise comparisons were conducted between the three groups from Project 1 and the three groups from Project 2. DEPs were identified based on |log₂(fold change)| > 1.2, and categorized as Log2FC_1, Log2FC_2, and Log2FC_3, as illustrated in Fig. 6f. The analysis showed that over 90% of DEPs were reproducibly measured across both projects. Furthermore, the Jaccard Similarity Index (JSI), a measure of quantitative similarity between datasets, was calculated for the three Log2FC groups, resulting in a value of 0.718. This high JSI score reflects excellent quantitative reproducibility of DEPs between the independent projects.

In addition, all the replicates of QC samples analyzed here were examined for potential contamination risks from platelets, erythrocytes, or coagulation, as highlighted by Geyer et al^15^. Quality marker panels across all replicates of the two projects were systematically checked, and none exhibited signs of obvious contamination (Supplementary Fig. S5, S6). Moreover, we compared the quantitative accuracy and fold-change accuracy between the reference neat digestion workflow and the Proteonano workflow, the results are Detailed results are presented in Supplementary Information, Fig. S7 and S8.

## Use of Proteonano workflow for neurodegenerative biomarker discovery

Emerging immunoassay-based and immuno-MS-based technologies have demonstrated excellent accuracy in detecting well-established blood-based biomarkers linked to cognitive decline, such as plasma Aβ, p-Tau, GFAP, NfL, and TDP-43^34,35^. However, these approaches are suitable for detecting a few validated biomarkers for the purpose of clinical diagnosis for a specific disease. Typically, they fall short in providing an unbiased discovery of novel plasma biomarkers, which could enable more nuanced patient stratification and facilitate early screening—not solely for Alzheimer’s pathology, but for neurodegenerative diseases in general.

Affinity-based multiplexed proteomics platforms like Olink and SomaLogic have been employed in the discovery of blood-based plasma biomarkers associated with cognitive decline^36^. However, these two approaches generally rely on panels of pre-selected proteins and are unable to differentiate post-translational modifications (PTMs), such as phosphorylation. In contrast, mass spectrometry offers an unbiased approach to profiling the plasma proteome and PTMs and has the potential to discover novel biomarkers. Nevertheless, it often faces challenges in detecting low-abundance blood-based biomarkers. Therefore, MS is traditionally applied to brain tissues and cerebrospinal fluids rather than plasma for neurological biomarker detection^37^. Thus, there remains a critical need for discovery tools that provide deep proteomic coverage, robust performance, and cost-effectiveness, addressing current gaps in biomarker identification.

We conducted a proof-of-concept study to demonstrate the feasibility of utilizing the Proteonano workflow for the discovery of potential plasma biomarker candidates. 183 plasma samples were collected from the Hubei Memory and Aging cohort Study (HAMCS) (ChiCTR; Registration number: ChiCTR1800019164), which included two tests for global cognitive function (the Mini-Mental State Examination [MMSE] and the basic Montreal Cognitive Assessment). The details of the population characteristics can be seen in the previous publications^38,39^ and Supplementary Table 3. In this experiment, a total of 195 plasma samples (Control: n = 125; Cognitive decline: n = 58; QC: n=12) were processed in six batches using the Proteonano workflow. The complete workflow was visualized in Fig. 7a.

**Figure 7.**
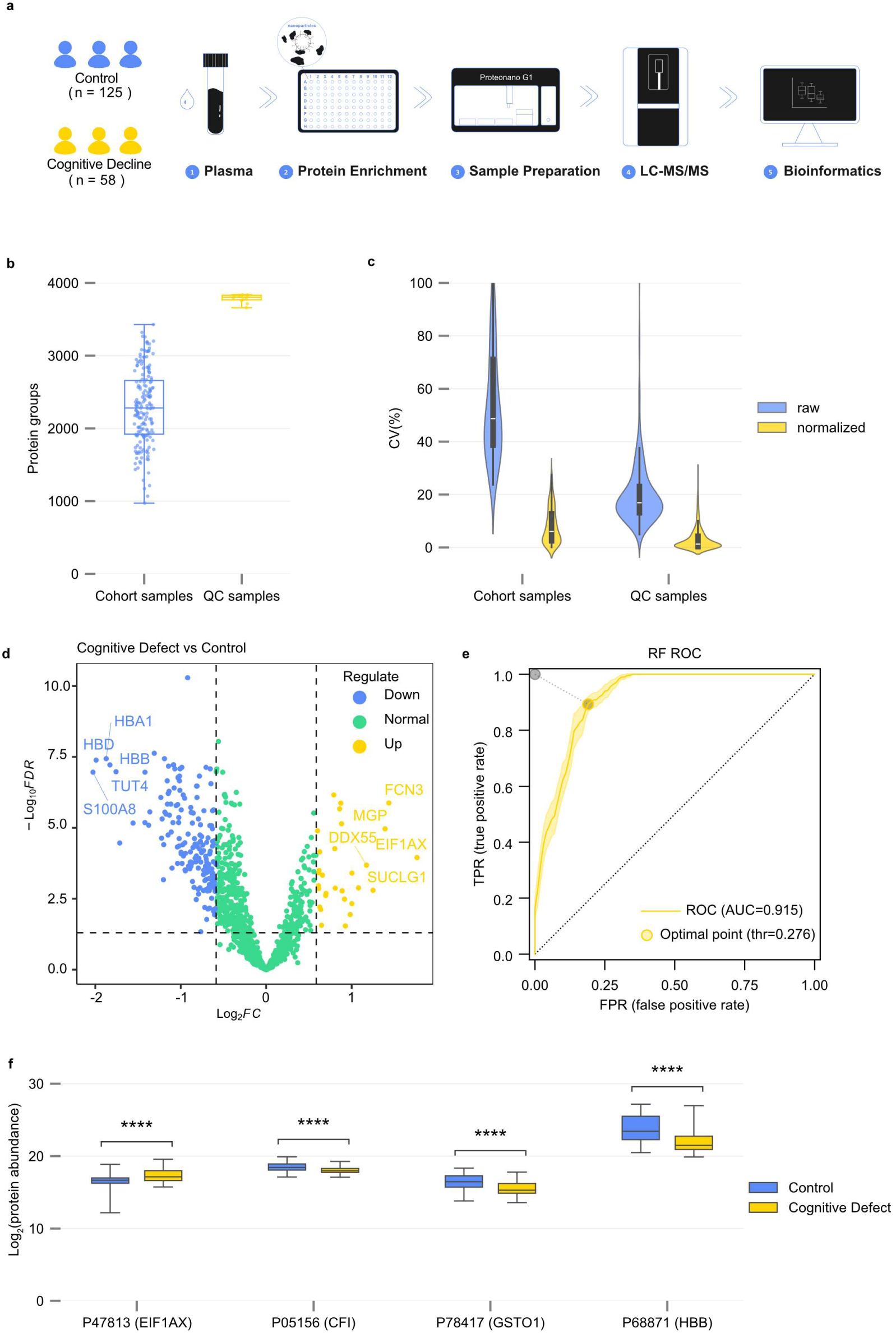
The feasibility of using the Proteonano workflow for the discovery of potential biomarker candidates. **a** Study design. **b** Number of proteins identified in the Cognitive Decline sample group and Control group. **c** Quantification precision assessed by calculating the CVs of the proteins in samples of the cognitive decline and control group. **d** Volcano plot of differentially expressed proteins in the control group and the cognitive decline group. **e** ROC curves of best multivariate model based on features selected by random forest method. **f** Illustration of selected proteins that are either increased (EIF1AX) or decreased (CFI, GSTO1, and HBB) in plasma with cognitive decline when compared with the ones in the Control Group.

We first assessed the quality of data obtained from 12 QC samples. As shown in Fig. 7b, an average of 3788 ± 16 protein groups (AVG±SE, n=12) were identified, with a median CV of 16.9%, indicating high quantitative consistency. For the cohort samples, an average of 2298 ± 38 protein groups (AVG±SE, n=183) were identified. However, the median CV for protein raw intensities prior to quantile normalization was **48.7%** (Fig. 7c), reflecting significant biological variability compared to technical variability. In their study on biomarker discovery for Parkinson’s disease via cerebrospinal fluid proteome profiling, Mann and colleagues^40^ highlighted that achieving technical variability lower than biological variability is crucial for identifying disease-specific proteomic signatures. This observation emphasizes the importance of producing high-quality and consistent data in biomarker research.

In addition, through differential protein expression analysis (Fig. 7d), we identified 8 upregulated proteins groups and 49 downregulated proteins groups (FDR corrected p value < 0.05 and |log2FC| > 1) between samples with and without cognitive function decline. Additionally, we matched these 57 DEPs with the abundance data from the HPPP database and revealed that 13 of these proteins fall within the low abundance range, defined as less than 10 ng mL^-1^. The DEPs are listed in Supplementary Table 4. For comparison, only 5 of the 13 low-abundance DEPs were detected in neat plasma without the use of Proteonano workflow. Further, we conducted a cross-examination to determine whether these 13 low abundance proteins have been detected by other plasma proteome profiling platforms. Notably, Soni et al.^41^ reported that 12 of the 13 DEPs were detectable using Seer’s Proteograph XT platform.

Furthermore, we found that several proteins among the DEPs have been previously linked to cognitive decline in earlier research, indicating their potential significance in neurodegenerative processes. We highlight seven proteins—GPX1, RAC1, RAC2, AZU1, S100A4, SUCLG1, and CDC42, all of which have been implicated in various neurodegenerative processes. GPX1, a selenium-dependent antioxidant enzyme, has shown protective effects against oxidative stress in neurodegenerative disorders^42^. RAC2, known for its role in cytoskeletal dynamics, regulates neurite outgrowth, and its function is impaired when γ-adducin cleavage leads to its downregulation^43^. CAP37 (AZU1), a protein released by neutrophils, promotes microglial activation and neuroinflammation in Alzheimer’s disease, with elevated levels in AD brains correlating with disease progression^44^. Similarly, S100A4 is associated with mitochondrial dysfunction and the accumulation of reactive oxygen species (ROS) in astrocytes affected by Alzheimer’s disease, contributing to neurodegenerative pathology^45^. SUCLG1, an essential enzyme in the TCA cycle, shows decreased modifications in Alzheimer’s disease; however, treatment with β-hydroxybutyrate has been shown to enhance its activity and ATP production, suggesting potential therapeutic avenues^46^. Lastly, CDC42, a key regulator of cell signaling, is involved in Alzheimer’s disease-related cognitive deficits and dendritic spine loss, with its decline directly correlating with disease progression and cognitive impairment^47,48^. These findings emphasize the significant roles of these proteins in neurodegenerative processes, highlighting them as potential targets and biomarkers for further research and therapeutic development.

Although many proteins require further investigation, we explored whether a small panel of proteins could reliably distinguish individuals with cognitive decline from those without. Differential protein expression data were analyzed using a random forest approach, which identified key features separating the two cohort groups. Based on these features, optimal multivariate models were constructed using the Akaike Information Criterion. The best model incorporated four proteins—EIF1AX, CFI, GSTO1, and HBB—and achieved an excellent AUC value of 0.92. Fig. 7e presents a receiver operating characteristic (ROC) curve, demonstrating the model’s ability to distinguish participants in the Cognitive Decline Group from the Control. Notably, GSTO1’s role in modulating oxidative stress stands out as a significant factor that may contribute to the underlying pathophysiological mechanisms of neurodegeneration^49^. This suggests its potential as a proteomic signature for patient stratification. However, transition from biomarker discovery to validation is a lengthy, time-consuming, and oftentimes expensive process. It involves critical steps of study design, including cohort selection, evaluation of statistical power, sample blinding, randomization, and rigorous data quality control^50–53^. While a detailed discussion of these best practices exceeds the scope of this study. Further efforts on the validation of their candidacy as potential biomarkers are necessary and will be the focus of our future work.

For cross-referencing, another nanoparticle-based MS proteomics platform, known as Proteograph XT, was utilized by Lucar et al.^54^ to analyze plasma samples from 379 AD patients and 240 healthy controls using a linear mixed-effects model to identify AD-associated proteins. Their findings revealed that proteins upregulated in AD included DKK2, MAPK3, HLA-DRA, LPL, and ACSL4, among others, while downregulated proteins included ETFA, TFRC, ADH1B, MGST3, and TNFRSF6B. Conversely, the Mag-Net enrichment strategy examined plasma samples from 10 AD patients and 10 healthy controls, applying a two-sided Mann-Whitney U test to identify AD-associated proteins^11^. This approach identified upregulated proteins in AD such as INPPL1, VPS13A, DNAJC10, S100B, and HTT, while downregulated proteins included VCP, MAP2K2, USP15, JMJD6, and CD36. Notably, our study observed a consistent trend in the differential expression of VCP with results reported by the Mag-Net strategy. However, none of the well-established blood biomarkers—such as Aβ, p-Tau, GFD, and TDP-42—were identified by any of those three nanoparticle-enriched mass spectrometry workflows. We hypothesize that factors like cohort characteristics, sample size, population diversity, and nanoparticle enrichment methodology may have contributed to this outcome. Further investigations with refined experimental designs are warranted to explore these observations more comprehensively.

## Discussion

This study developed and systematically evaluated the Proteonano Ultraplex Platform for robust, deep plasma proteomic analysis. When paired with advanced mass spectrometers, the platform demonstrated exceptional capability for deep proteomic analysis of low-input plasma samples, achieving extensive proteome coverage, high throughput, a dynamic range spanning nine orders of magnitude, and minimized batch-to-batch variability. Furthermore, we demonstrated that the Proteonano workflow exhibited remarkable stability and reproducibility across ten distinct and widely adopted DIA LC-MS/MS systems, showcasing flexibility in balancing protein depth and sample processing speed. Finally, the platform’s performance was validated using plasma samples from an elderly community cohort, including individuals with neurodegenerative diseases.

The PUP Platform described here offers several advantages for MS-based plasma proteomics studies, especially in the discovery phase of biomarker screening. Unlike traditional immunodepletion methods, our platform is highly standardized, robust, high-throughput, user-friendly, cost-effective, and compatible with various mass spectrometry configurations across different laboratories. Notably, the workflow requires only 40 µL of plasma per sample to achieve optimal performance, whereas previously reported technologies typically require at least 100 µL^12^. This critical feature, which has not been emphasized enough, plays a key role in the practical adoption of new technologies, especially given the common limitations on sample volumes. In comparison, Olink PEA technology, widely adopted by dozens of biopharmaceutical companies and applied to millions of samples globally, requires only 10 µL of plasma.

Secondly, the PUP Platform achieved cutting-edge deep plasma proteome coverage and data quality by effectively combining multiple nanoparticles into a single, uniform reagent, a strategy akin to those utilized in multiplexed immunoassays but without the challenges of antibody cross-reactions. This innovative approach removes the need for sample fractionation and multiple mass spectrometry runs for individual samples. As a result, it significantly minimizes time, cost, and batch effect in plasma proteomics studies, making it particularly advantageous for large cohorts involving hundreds or even thousands of samples. In the future, we plan to enhance nanoparticle technology by integrating a wider range of compositions into a single reagent to enrich a broader range of low abundance plasma proteins. This innovation aims to achieve deeper plasma proteome coverage through MS-based proteomics, bringing its capabilities closer to those of non-MS multiplex technologies regarding proteome coverage, such as SomaScan 7K and 11K^55^.

Thirdly, the PUP Platform has implemented a fully standardized workflow, starting with nanoparticle-plasma incubation at the beginning of the experiment and extending to data analysis across various batches. This carefully structured system incorporates a robust quality control framework, guarantees compatibility with multiple mainstream mass spectrometers, supports deep plasma coverage, and minimizes batch-to-batch effect over independent studies. This level of standardization significantly improves reliability and efficiency, solidifying its applicability to large-scale plasma proteomics studies.

That said, we have not included discussions of other considerations that may be associated with the proteome coverage and data reproducibility of nanoparticle-enriched plasma proteomics. For instance, the effect of nanoparticle size and size distribution on the compositions of protein corona is less studied. Notably, Tenzer et al.^23^ demonstrated the significance of nanoparticle size in shaping corona layers by using monodispersed silica nanoparticles of varying sizes (20, 30, and 100 nm). Their findings revealed that nanoparticle size critically influenced the composition of the corona, which accounted for 37% of the identified proteins in their study^23^. Yet, the effect of size uniformity on the coverage and the reproducibility of proteomic analysis remains unexamined. In this context, the surface-modified nanoparticles presented here offer an average diameter of 300 nm and exhibit exceptional size uniformity (PDI < 0.05, Fig. 3b). In comparison, particles documented by Blume^10^ and Wu^11^ are either characterized by relatively high polydispersity or, in the case of microparticles, lack detailed data on size distribution. Alternative approaches, such as graphene oxide-modified materials, have also been explored for proteome enrichment^56^. However, the highly heterogeneous nature of these flake-like nanomaterials may pose substantial challenges on quality control, particularly when employed in large cohort plasma studies. Although the precise mechanisms remain unclear, we hypothesize that employing nanoparticles with defined size and low polydispersity is crucial for ensuring quality control and enhancing reproducibility in plasma proteomics applications^57^.

In addition, we recognize that the proof-of-concept study using the PUP Platform for biomarker discovery is subject to several limitations. First, mass spectrometers inherently face significant challenges in detecting low-abundance proteins in plasma. This limitation is particularly evident for biomarkers of neuropathology, which are present at very low concentrations in plasma (typically a few pg.mL^-1^). That explains partly why none of the three reported nanoparticle-enriched and MS-based workflows detected low abundant biomarkers like p-Tau181. We cannot rule out the possibility that these AD-related biomarkers were also present in this cohort but at concentrations below the detection sensitivity of mass spectrometry, even after enrichment. While the PrecivityAD2 assay also relies on mass spectrometry, it incorporates the use of antibody-conjugated microbeads to enrich specific AD biomarkers from plasma prior to LC-MS/MS measurements. Second, it is evident that the cohort is relatively small and lacks diversity. Mann and colleagues proposed a “rectangular” plasma proteome profiling strategy that suggested ^29^ typically 100 to 1000 samples should be used in the phase of plasma biomarker discovery.

Therefore, we anticipate that PUP Platform could serve as a valuable discovery-phase tool, enabling the profiling of thousands of plasma proteins to identify potential biomarkers. These identified biomarkers can then undergo further validation in larger cohorts using targeted proteomics techniques such as MS-based Parallel Reaction Monitoring (PRM) and single-molecule immunoassays like Simoa, or even low-cost ELISA.

In summary, our study highlights the Proteonano Ultraplex Proteomics Platform as an innovative solution for MS-based deep plasma proteomics. This platform offers extensive proteomic coverage, high-throughput capabilities, cost efficiency, and a standardized data production workflow. It supports minimal plasma input (40 µL), demonstrates compatibility with various LC-MS/MS systems and throughput levels, and ensures batch-to-batch consistency, reproducibility, and precision in protein identification and analysis in large-cohort studies. These advancements offer exceptional potential to push the frontiers of plasma proteomics and pave the way for the discovery of novel protein biomarkers and therapeutic targets across a range of diseases. In future endeavors, we aim to expand our research by evaluating the performance of the PUP platform across multiple laboratories and conducting larger, multi-centered plasma proteomics studies to further validate its robustness and scalability.

## Methods

### Blood Samples

For most of the experiments, pooled healthy donor blood samples were obtained from Oricells (Ori Biotech, Shanghai, China), Whole blood was collected in K2-EDTA containing tubes. Samples were centrifuged at 1500 *g* for 15 min at 4 °C. Supernatant was transferred from centrifuge tubes. Plasma samples from multiple donors were pooled, aliquoted into microfuge tubes, and stored at −80 °C until analysis.

Plasma samples collected from a community cohort aimed at understanding cognitive deficiencies in elderly patients in Hubei Province (Hubei Memory and Aging Cohort Study, HMACS) were utilized. Ethical approval for blood sample collection was received at respective institutions for the study.

## Preparation and Characterization of Nanoparticles

The magnetic Fe_3_O_4_ particles were prepared according to a solvothermal approach. Typically, FeCl_3_·6H2O (3.91 g) and trisodium citrate (0.46 g) were first dissolved in ethylene glycol (70 mL). Then, NaAc (9.42 g) was added, and the mixture was vigorously stirred to form a transparent solution. Afterward, the solution was transferred to a 150 mL Teflon-lined stainless-steel autoclave. The autoclave was sealed and heated at 200 °C and maintained for 12 h, and then it was allowed to cool to room temperature. The products were washed with ethanol and deionized water several times and dried at 60 °C for 12 h.

In order to prepare monodispersed Fe_3_O_4_@PDA (polydopamine) core-shell nanospheres, 180 mg of the hydrochloride dopamine was dispersed in 90 ml Boric acid-borax buffer (0.5 mol.L^-1^, pH=8.0). Then, 90 mg of the as-prepared Fe_3_O_4_ was added, stirring overnight at room temperature. The resultant product was separated and collected with a magnet and then washed with ultrapure water three times.

In a typical preparation of Fe_3_O_4_@PDA@Au, firstly, 20 mg of the Fe_3_O_4_@PDA was added into 12.5 mL deionized water, sonicate for 10 min. Then, 0.5 mL dopamine solution (0.76 mg of the hydrochloride dopamine was dispersed in 1 mL deionized water) and 83.5 μL sodium hydroxide standard solution (0.6M) were added in sequence. Typically, 0.8 mL 1 wt% aqueous solution of HAuCl_4_ (1 g HAuCl_4_ was dispersed in 100 mL deionized water) was quickly added and reaction was allowed for overnight at room temperature under shaking at 1000 rpm. Finally, the product was separated by a magnet and washed with water.

The nanomaterial sample was ultrasonically dispersed in ethanol, and a drop of the solution was then placed onto an ultra-thin carbon film. After drying, the sample was analyzed using a transmission electron microscope (TEM). The instrument used was an FEI Tecnai G2 F20 with a maximum acceleration voltage of 200 kV. It has a point resolution of 0.24 nm, a line resolution of 0.102 nm, and an information resolution of ≤0.14 nm. Energy dispersive spectroscopy (EDS) was performed using an Oxford XPLORE system.

Well dispersed Fe_3_O_4_ microspheres were prepared by a solvothermal method^58^. Fe_3_O_4_ microspheres were dispersed in chloroauric acid containing sodium citrate aqueous solution and stirred at 90 °C for 0.5 h. Resulting Au NPs-deposited Fe_3_O_4_ (Fe_3_O_4_@PDA@Au) were magnetically separated from the suspension and subsequently washed dried under vacuum.

For peptide conjugation, peptides were dissolved in water. Dissolved peptide was incubated with Tris(2-carboxyethyl) phosphine hydrochloride (TCEP) solution, followed by adding of Fe_3_O_4_@PDA@Au synthesized above. The reaction mixture was incubated at 25°C overnight. Reaction product was washed by ethanol then deionized water and stored at refrigerator at 4°C. Three peptides, PP1 (HKAATKIQASFRGHITRKKLC), PP2 (DIEEVEVRSKYFKKNERTVEC), and PP3 (DIEEVEVRSKYFKKNERTVEC) were custom synthesized by GenScript (Suzhou, China). The details can be seen in the Supporting Information S1 section.

Following peptide conjugated Fe_3_O_4_@PDA@Au synthesis, transmission electron microscopy (TEM) was performed by using a Tecnai 12 electron microscope (ThermoFisher Scientific) at an accelerating voltage of 200 kV. DLS and Zeta potential of the particles were examined by using a NanoBrook 90Plus PALS instrument (Brookhaven Instruments, Holtsville, NY).

## Peptide Conjugation Density

First, 125 μL of TCEP solution (200 mM) and 5 mL of FITC-peptide solution (0.2 mg.mL^-1^) were added to a centrifuge tube, mixed thoroughly, and incubated in the dark for 2 hours. Then, varying volumes (0, 200, 400, 600, and 1200 μL) of the TCEP-treated FITC-peptide solution (fluorescence intensity denoted as I₀) were transferred into separate 5 mL centrifuge tubes. Each tube was supplemented with 1000 μL of Fe₃O₄@PDA@Au NPs (1.5 mg.mL^-1^). Ultra-pure water was then added to bring the final volume to 2300 μL, followed by ultrasonic mixing for 5 minutes. The reaction mixtures were incubated at 25°C under shaking conditions for 17 hours. After the reaction, the samples were washed three times with ultra-pure water, and the supernatants from the three washes were collected (fluorescence intensity denoted as I₁). All fluorescence intensity values were measured in triplicate using a Tecan SPARK multifunctional microplate reader.

## Sample Preparation by the Proteonano Workflow

Automatic sample preparation was performed in batches utilizing Nanomation G1 workstation, equipped with both a magnetic module and a heater shaker module. Plasma samples were dispensed on 96 well flat bottom assay plates for subsequent processing. For most experiments, 40 μL of human plasma was diluted to a final volume of 100 μL by using 1 x PBS at pH7.4 and was subsequently combined with PCNPs (as synthesized above) from the Proteonano reagent in a 1:1 volumetric ratio. The mixture was incubated at 25 °C and agitated for 60 min. Following magnetic immobilization, PCNPs were washed thrice with 1 x PBS. Proteins captured on the PCNPs were reduced by 20 mM DTT at 37 °C for 60 min. Alkylation was performed using 50 mM iodoacetamide (IAA) at room temperature in darkness for 30 min. Trypsin (Promega Corporation, Madison, WI, USA) digestion was carried out at 37°C for a duration of 16 hours with shaking at 1000 rpm. Post-digestion, peptides were purified using desalting C18 columns in micropipette tip format (ThermoFisher Scientific) and lyophilized with a LyoQuest freeze dryer (LyoQuest, Telstar, Terrassa, Spain). Lyophilized peptides were then reconstituted in 0.1 % formic acid prior to mass spectrometry. Peptide concentrations were measured with a Nano300 microvolume spectrophotometer (Allsheng Instruments, China).

## LC-MS/MS Experiments

Multiple LC-MS/MS instruments were used in the study. For most studies, Orbitrap Astral (ThermoFisher Scientific) mass spectrometer was used. In some cases, Orbitrap Exploris 480 (ThermoFisher Scientific) mass spectrometer equipped with FAIMS, or timsTOF Pro 2 mass spectrometer (Bruker Instruments) were used.

For studies using Orbitrap Astral instrument, 300 ng or 500 ng peptides dissolved in 0.1% formic acid were separated by a Vanquish™ Neo UHPLC system (ThermoFisher Scientific) followed by mass spectrometry. Data were acquired in data independent acquisition mode (Detailed experimental setup are listed in Supplementary Table S5-14).

For studies using Orbitrap Exploris 480, 500 ng of peptides were separated by an Easy-nLC1200 reverse-phase HPLC system (ThermoFisher Scientific) using a precolumn (homemade, 0.075 mm × 2 cm, 1.9 μm, C18) and a self-packed analytical column (0.075 mm × 20 cm, 1.9 μm, C18) over a 48 min gradient before nano-electrospray on Orbitrap Exploris 480 mass spectrometer equipped with FAIMS (ThermoFisher Scientific). Solvent A was 0.1 % formic acid and solvent B was 80 % acetonitrile (ACN)/0.1 % formic acid. Gradient conditions were 3–7 % B (0–1 min), 7–30 % B (1–36 min), 30–95 % B (36–38 min), and 95 % B (38–48 min). Mass spectrometer was operated in DIA mode. Spray voltage was set to 2.4 kV, RF Lens level at 40 %, and heated capillary temperature at 320 °C. Full MS resolutions were set to 60,000 at m/z 200 and full MS automatic gain control (AGC) target was 100 % with an injection time (IT) of 50 ms. Mass range was set to m/z 350–1200. The AGC target value for fragment spectra was set at 1,000 %. Resolution was set to 30,000 and IT to 54 ms. Normalized collision energy was set at 30 %. Default settings were used for FAIMS with voltages applied as -45 V, except gas flow, which was applied with 3.5 L/min.

For studies using timsTOF Pro 2 instrument, 200 ng of peptides was subjected to a nanoElute® 2 nanoLC system coupled with a Bruker timsTOF Pro 2 mass spectrometer using a trap-and-elute configuration. First, the peptides were loaded on an Acclaim PepMap 100 C18 trap column (0.3 mm ID × 5 mm), then separated on a self-packed analytical column (0.075 mm × 25 cm, 1.8 μm, C18) at a flow rate of 300 nL min^−1^. Solvent A was 0.1 % formic acid and solvent B was 80 % ACN/0.1 % formic acid. Gradient conditions were 2–22 %

B (0–45 min), 22–35 % B (45–50 min), 35–80 % B (50–55 min), 80 % B (55–60 min). The spray voltage was set to 1.5 kV. The mass spectrometer was operated in diaPASEF mode using ion mobility range of 0.76–1.29 Vs/cm^2^ with 100 ms accumulation time. MS1 mass spectrometry scans 452-1152 m/z with peak height above 2,500 before being detected. The 452-1152 m/z range was divided into four steps. Each step was divided into seven windows, the Number of Mobility Windows was set to 2, and a total of 56 windows for continuous window shattering and information gathering. The splitting mode was CID and the splitting energy was set 20-59 eV. The mass width of each window was 25 Da and the cycle time for a DIA scan was 1.59 s.

## Proteomic Data Analysis

The *.raw* or *.d* files were obtained directly from mass spectrometers, or *mzML* files converted from .raw files by msConvert^59^ (Version 3.0) software were searched using DIA-NN (Version 1.8.1) in library free mode. For each sample, spectra were searched against a UniProt *Homo sapiens* reviewed proteome dataset, or other datasets. DIA-NN search parameters were: 10 ppm mass tolerance for mass accuracy, one missed cleavages of trypsin, carbamidomethylation of cysteine as fixed modification, and methionine oxidation as the only variable modification. The rest of the parameters were set to default. The FDR cutoffs at both precursor and protein level were set to 0.01.

The .raw or .d files generated directly from mass spectrometers, or mzML files converted from .raw files using msConvert (Version 3.0) software, were analyzed with DIA-NN (Version 1.8.1) in library-free mode. For each sample, spectra were matched against a UniProt Homo sapiens reviewed proteome dataset or other relevant datasets. The DIA-NN search parameters included a 10 ppm mass tolerance for mass accuracy, one missed cleavage for trypsin, carbamidomethylation of cysteine as a fixed modification, and methionine oxidation as the sole variable modification. All other parameters were set to default, with false discovery rate (FDR) cutoffs of 0.01 applied at both the precursor and protein levels.

## Statistical Analysis and Data Visualization

Coefficient of variation (CV) for each protein was determined by dividing its empirical standard deviation by its empirical mean, and CV analyses were performed on raw intensities and quantile-normalized intensities, respectively. Median values were reported as overall coefficient of variation. Pearson correlation analyses and linear regression were conducted using Pingouin^59^. No missing value imputation was performed during the above analysis, and calculations were performed only for proteins that were identified in all samples. For replicate experiments, protein identifications are expressed as AVG±SE. Seaborn was used to generate bar charts and violin charts^60^. The matplotlib-venn^61^ package (https://pypi.org/project/matplotlib-venn/) and the Venn^62^ (https://pypi.org/project/venn/) package were used to plot VENN^39^ plots. MS-DAP was used to screen for differentially expressed proteins, and ggVolcano^63^ (https://github.com/BioSenior/ggVolcano) was used to generate volcano plot. KEGG pathway enrichment analysis was performed using the GSEApy^64^ software package in Python. Feature selection was performed using MetaboAnalystR^65^. Multivariate analysis was performed using MASS^66^ in R. Variables were first processed by glm (MASS package in R), followed by Akaike information criterion (AIC) determination using stepAIC (MASS package in R) with forward selection and Wald test. Multivariate Receiver operating characteristic(ROC) analysis was subsequently performed using roc (glmtoolbox^67^ package in R) to assess the predictive performance of the logistic regression model selected by step AIC. ROC curves were graphed using ROC-utils^68^.

## Data Availability

All raw and processed proteomics data generated in this study have been deposited to the iProX repository and are available for review. The data can be accessed at https://www.iprox.cn/page/PSV023.html?url=1744530654097Tmnc using the password dO25. The data are currently under controlled access for peer review and will become publicly available after the embargo period. Alternatively, the data can be made available upon reasonable request by contacting the corresponding author.

## Acknowledgements

This work was financially supported by Nanomics Biotechnology., Ltd.

## Author contributions

H.W. designed the study and drafted the manuscript. Y.W. led product development. Y.H.Z. performed the proteomics data analysis. X.H.O. conducted the sample preparation and mass spectrometry experiments.

