## Supplemental file for "Proteonano™: A Robust Platform for Large Cohort Deep Plasma Proteomic Studies"

### 1. Proteonano nanoparticles

The Proteonano Enrich Kit is composed of hierarchically structured nanoparticles functionalized with peptides (PCNPs). The physicochemical properties of three peptides used in this work are shown in Supplemental Table 1.

**Supplemental Table 1.** The physicochemical properties of three peptides

| Name | Sequence | Length | $M_w$ | Isoelectric Point | Charge (mV) | Hydrophobicity | GRAVY |
|---|---|---|---|---|---|---|---|
| PP1 | HKAATKIQASFRGHITRKKLC | 21 | 2,395 | 11.73 | 0.30 | 38% | -0.65 |
| PP2 | DIEEVEVRSKYFKKNERTVEC | 21 | 2,602 | 4.90 | -1.04 | 62% | -1.22 |
| PP3 | QETLKDTRSKFFNKPSMTVVC | 21 | 2,460 | 9.73 | 1.95 | 48% | -0.63 |

### 2. Layout of the Nanomation workstation

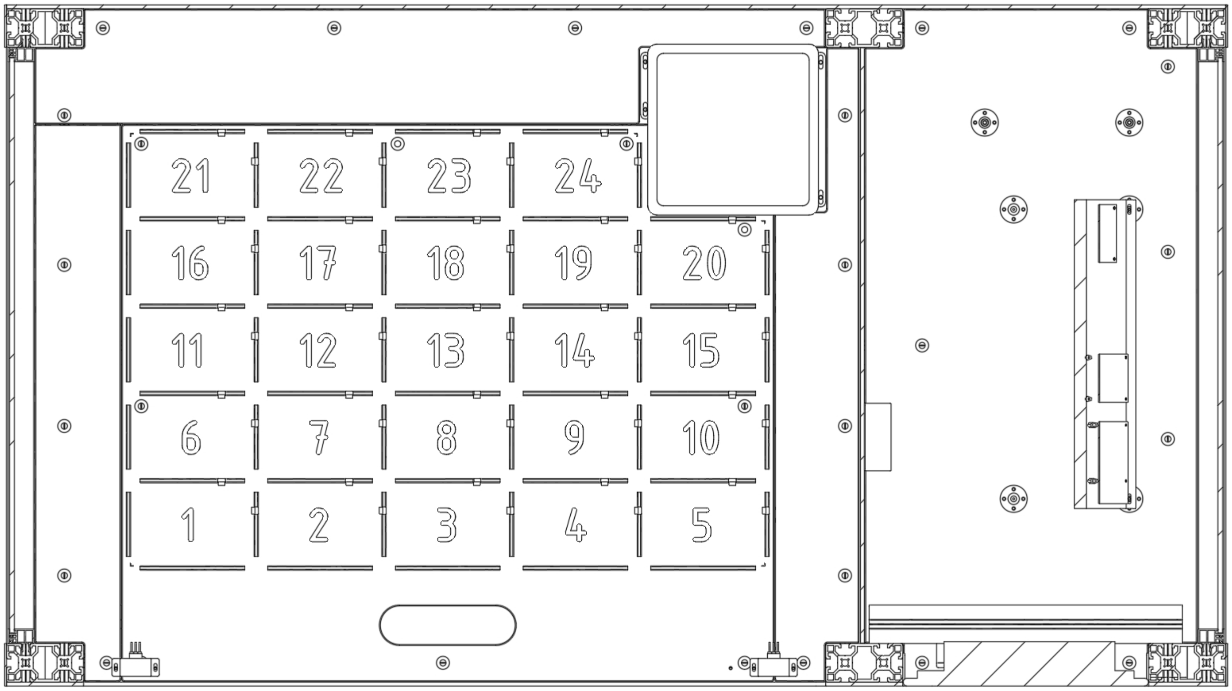

**Supplemental Figure. 1** The top view of the Nanomation workstation.

### 3. Quality control sample configuration

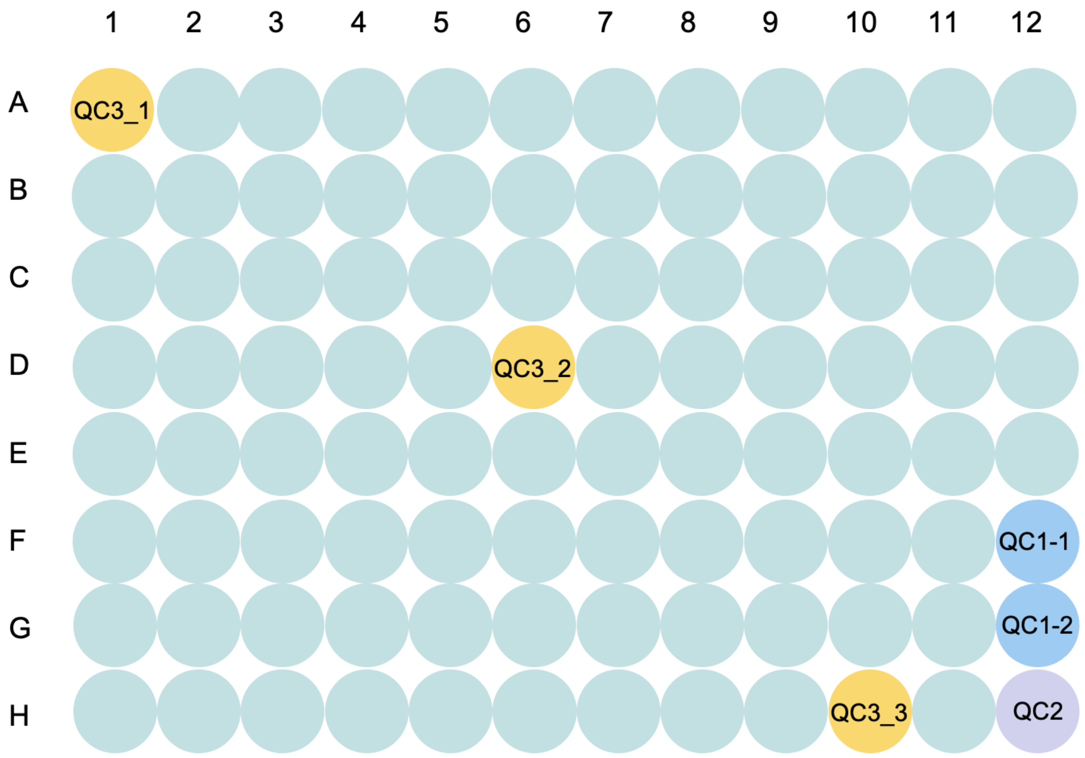

**Supplemental Figure. 2** A typical quality control sample configuration on a 96 well plate. **QC 1**: QC1 is a lyophilized peptide mix derived from pooled healthy human plasma that has undergone protein enrichment, reduction, alkylation, enzymatic digestion, and desalting. QC1 is used to monitor the reproducibility of peptide signal readout by LC-MS/MS instruments and typically performed twice on a 96 well plate. **QC 2**: QC2 uses pooled healthy donor plasma without being enriched (neat plasma) and processed to undergo sample reduction, alkylation, enzymatic digestion, desalting, and lyophilization, thus monitoring the quality of conventional procedures during sample preparation. Usually, one QC2 sample is included per 96-well plate. **QC 3**: QC3 uses pooled healthy donor plasma, but undergoes nanoparticle-based protein capture, in addition to conventional steps of MS-based proteomic sample preparation. Thus, the performance of the complete processing pipeline is monitored. By comparing results obtained from QC1 and QC2, the performance of the protein enrichment process can be deducted. Three replicates of QC1 are included in each fully loaded 96-well plate.

### 4. Specifications of ten LC-MS/MS systems

**Supplemental Table 2.** Specifications of ten LC-MS/MS systems.

| LC-MS System | LC Column | Mass Spectrometer | Throughput |
|---|---|---|---|
| 1 | Easy-nLC1200 | Orbitrap Exploris 480 | 24 SPD |
| 2 | nanoElute 2 | timsTOF Pro 2 | 24 SPD |
| 3 | ES906 | Orbitrap Astral | 180 SPD |
| 4 | ES906 | Orbitrap Astral | 100 SPD |
| 5 | ES906 | Orbitrap Astral | 60 SPD |
| 6 | ES906 | Orbitrap Astral | 24 SPD |
| 7 | ES75550 | Orbitrap Astral | 14 SPD |
| 8 | µPac 110 cm | Orbitrap Astral | 15 SPD |
| 9 | µPac 110 cm | Orbitrap Astral | 11 SPD |
| 10 | µPac 110 cm | Orbitrap Astral | 7 SPD |

### 5. Batch-to-batch reproducibility of Proteonano workflow

To assess the reproducibility of the Proteonano workflow, two independent batches of analysis were performed on eight samples: A1–A4 (four replicates from a healthy male donor) and E1–E4 (four replicates from a healthy female donor). Pearson correlation coefficients were calculated to evaluate consistency across samples. For Group A, the mean correlation coefficients were 0.986 and 0.983 in the first and second batches, respectively. For Group E, the mean correlation coefficients were 0.994 and 0.992 in the respective batches, highlighting the high reproducibility of the workflow.

a

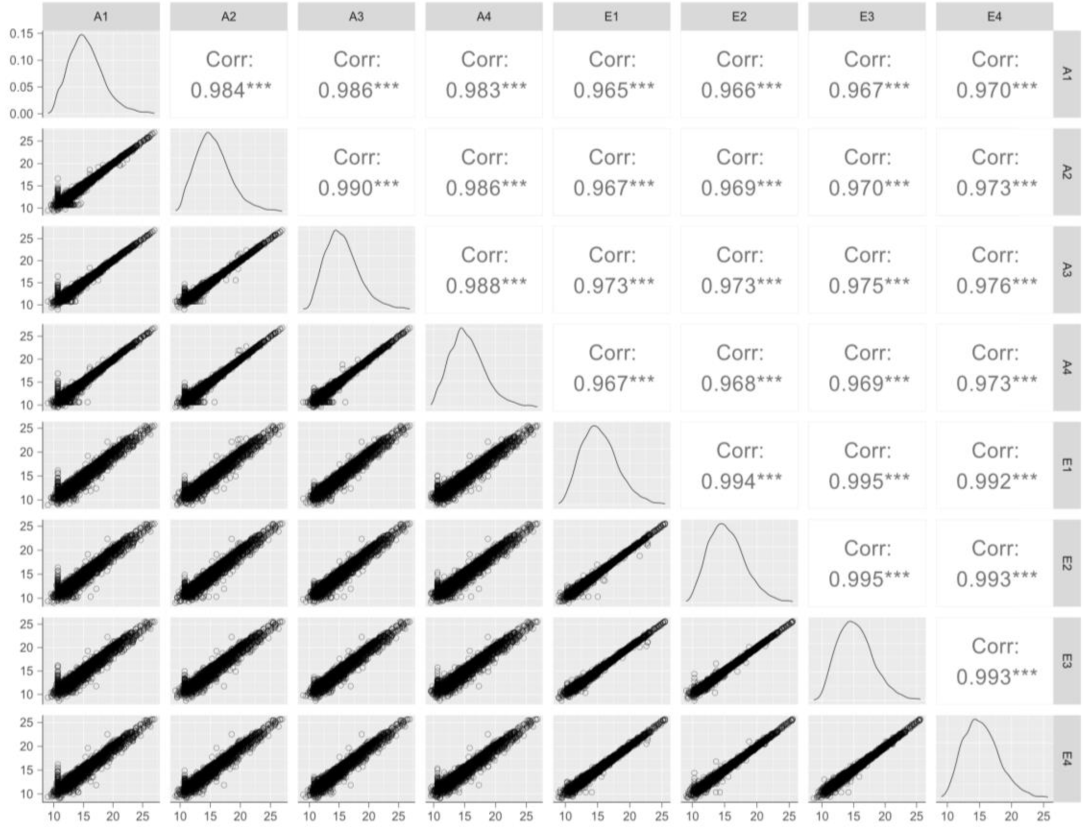
A1
A2
A3
A4
E1
E2
E3
E4
Corr: 0.984***
Corr: 0.986***
Corr: 0.983***
Corr: 0.965***
Corr: 0.966***
Corr: 0.967***
Corr: 0.970***
Corr: 0.990***
Corr: 0.986***
Corr: 0.967***
Corr: 0.969***
Corr: 0.970***
Corr: 0.973***
Corr: 0.988***
Corr: 0.973***
Corr: 0.973***
Corr: 0.975***
Corr: 0.976***
Corr: 0.967***
Corr: 0.968***
Corr: 0.969***
Corr: 0.973***
Corr: 0.994***
Corr: 0.995***
Corr: 0.992***
Corr: 0.995***
Corr: 0.993***
Corr: 0.993***

b

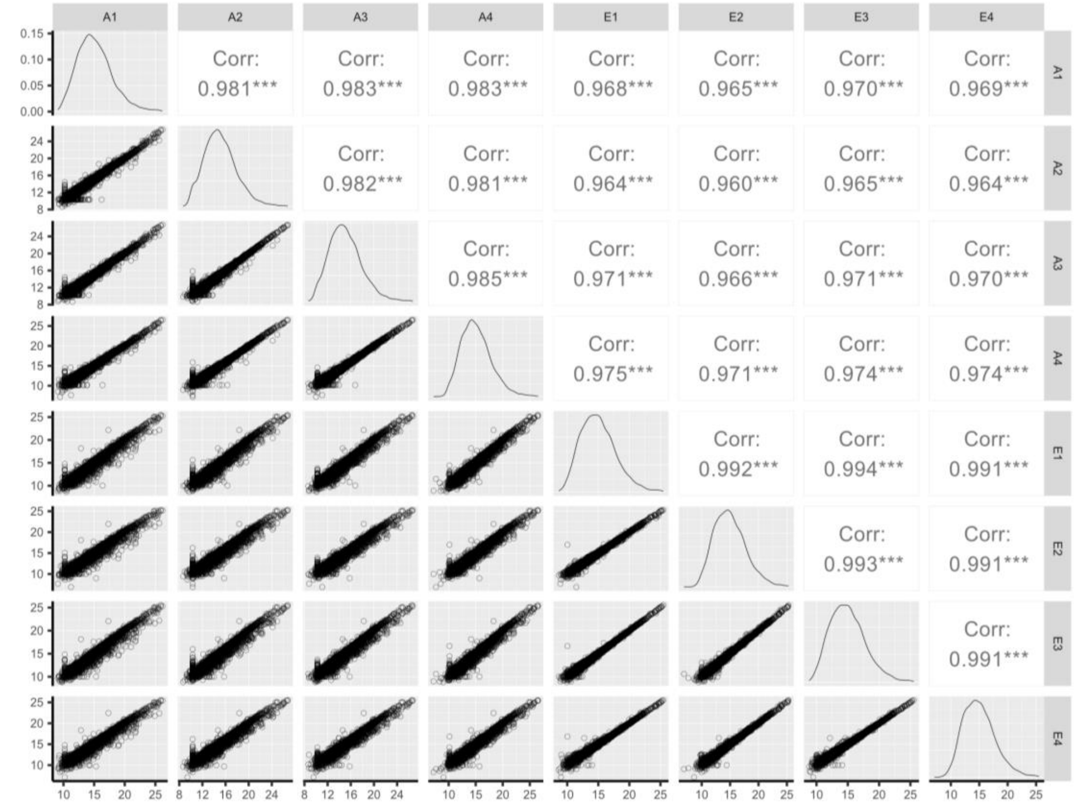
A1
A2
A3
A4
E1
E2
E3
E4
Corr: 0.981***
Corr: 0.983***
Corr: 0.983***
Corr: 0.968***
Corr: 0.965***
Corr: 0.970***
Corr: 0.969***
Corr: 0.982***
Corr: 0.981***
Corr: 0.964***
Corr: 0.960***
Corr: 0.965***
Corr: 0.964***
Corr: 0.985***
Corr: 0.971***
Corr: 0.966***
Corr: 0.971***
Corr: 0.970***
Corr: 0.975***
Corr: 0.971***
Corr: 0.974***
Corr: 0.974***
Corr: 0.992***
Corr: 0.994***
Corr: 0.991***
Corr: 0.993***
Corr: 0.991***
Corr: 0.991***

**Supplemental Figure. 3 Pairwise correlation of protein intensities among samples within each batch. a** Batch 1 and **b** Batch 2. Each point represents the intensity of a protein in a pair of samples.

We next tested reproducibility of sample preparation through the Proteonano workflow using the PCNPs produced in two separate batches. This allowed us to test both variations introduced by plasma sample processing and nanoparticle synthesis. Aliquots of the same pooled human plasma sample were parallelly processed using the Proteonano workflow, followed by LC-MS/MS measurements. The number of protein groups identified in Proteonano workflow processed plasma samples for PCNPs synthesized in batch 1 was 4057 ± 38 (AVG ± SE, n=12), with a CV of 9.87 % on the raw intensity and a CV of 4.07 % on the normalized intensity, while protein groups detected for PCNPs synthesized in batch 2 was 4159 ± 30 (AVG ± SE, n=11), with a CV of 9.62 % on the raw intensity and a CV of 4.03 % on the normalized intensity (Supplementary Fig. 4a-c). Venn diagram further demonstrated that similar protein groups were identified in both batches of PCNPs (Supplementary Fig. 4b). To assess the correlation of relative abundance of detected proteins among all MS runs, pairwise Pearson correlation coefficients were determined. The minimum correlation coefficient was 0.981 for PNCPs of batch 1, and 0.956 for PCNPs of batch 2, and the minimum correlation coefficient in all MS runs for both batches was 0.857, with a median of 0.889 (Supplementary Fig. 4d), demonstrating relatively excellent correlation of the two batches of PCNPs going through the complete enrichment-based plasma proteomics workflows

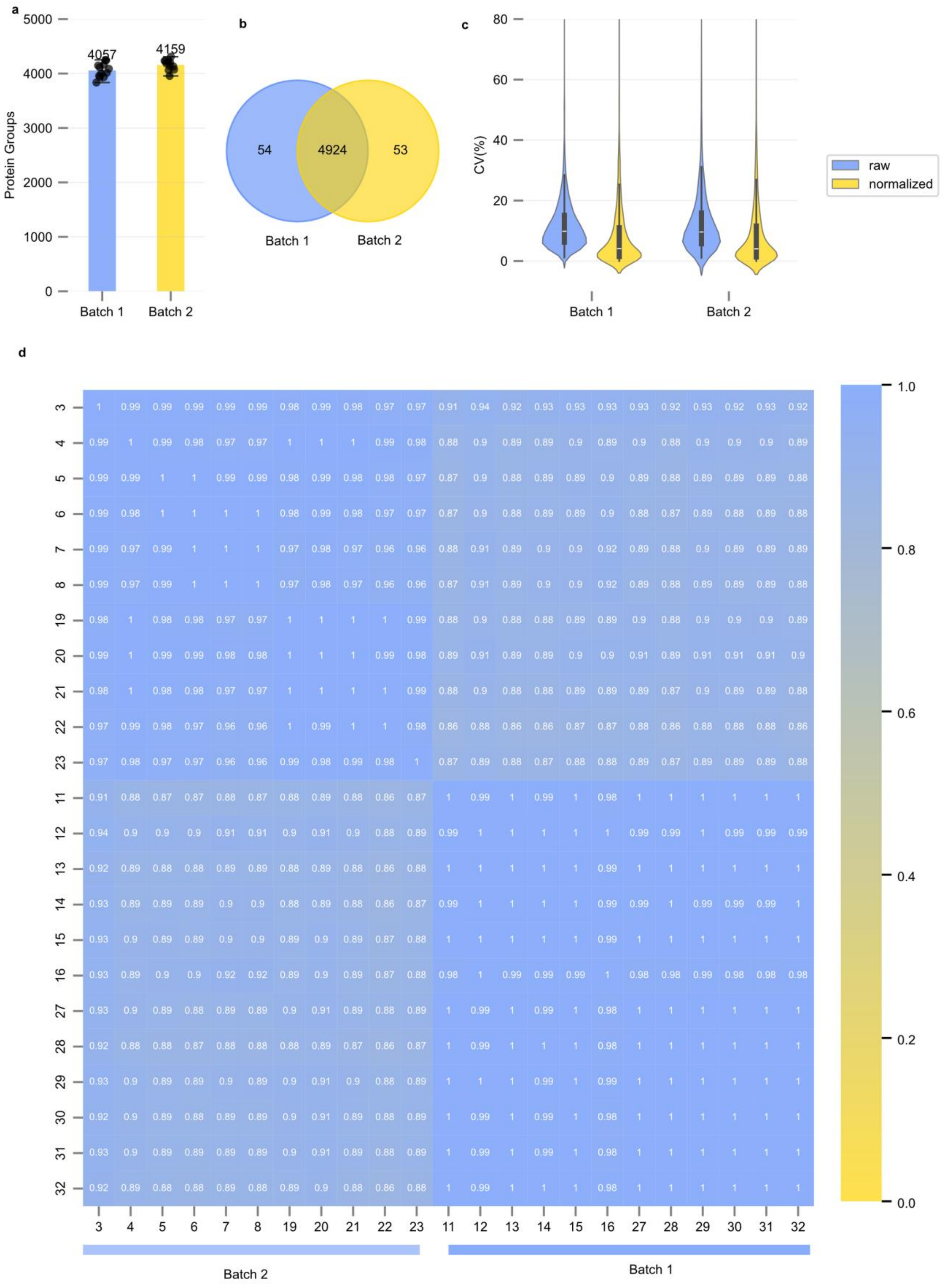
a
Protein Groups
5000
4000
3000
2000
1000
0
4057
4159
Batch 1
Batch 2
b
54
4924
53
Batch 1
Batch 2
c
CV(%)
80
60
40
20
0
raw
normalized
Batch 1
Batch 2
d
3 4 5 6 7 8 19 20 21 22 23 11 12 13 14 15 16 27 28 29 30 31 32
1.0
0.8
0.6
0.4
0.2
0.0
Batch 2
Batch 1

**Supplementary Figure 4. Proteonano workflow affords reproducibility over replicate batches of nanoparticles. a** Number of protein groups identified in each sample injection for samples processed using PCNPs from synthesis batch 1 and batch 2. Bar height and numbers represent the mean number of protein groups detected in each sample group. Individual dots correspond to protein groups detected in each MS run. **b** Venn diagram showing the overlap of protein groups identified in batch 1 and batch 2. **c** Quantification precision assessed by calculating the intra-plate coefficient of variation for all protein groups from the raw and normalized intensity matrices, based on repeated processing of the same sample using two batches of synthesized PCNPs. **d** Pearson correlation coefficients between samples.

### 6. Assessment of sample contamination

In order to assess possible platelet, erythrocytes, and coagulation contamination during plasma collection, Based on definitions established in previous studies[1], we evaluated the quality of each sample individually by calculating three contamination indices and examining their distributions. For each index, samples with values exceeding two standard deviations above the mean were initially identified as potentially contaminated (red lines in the Supplemental Fig. 5, Supplemental Fig. 6)**,** indicating that there is no significant contamination in these samples in the two projects.

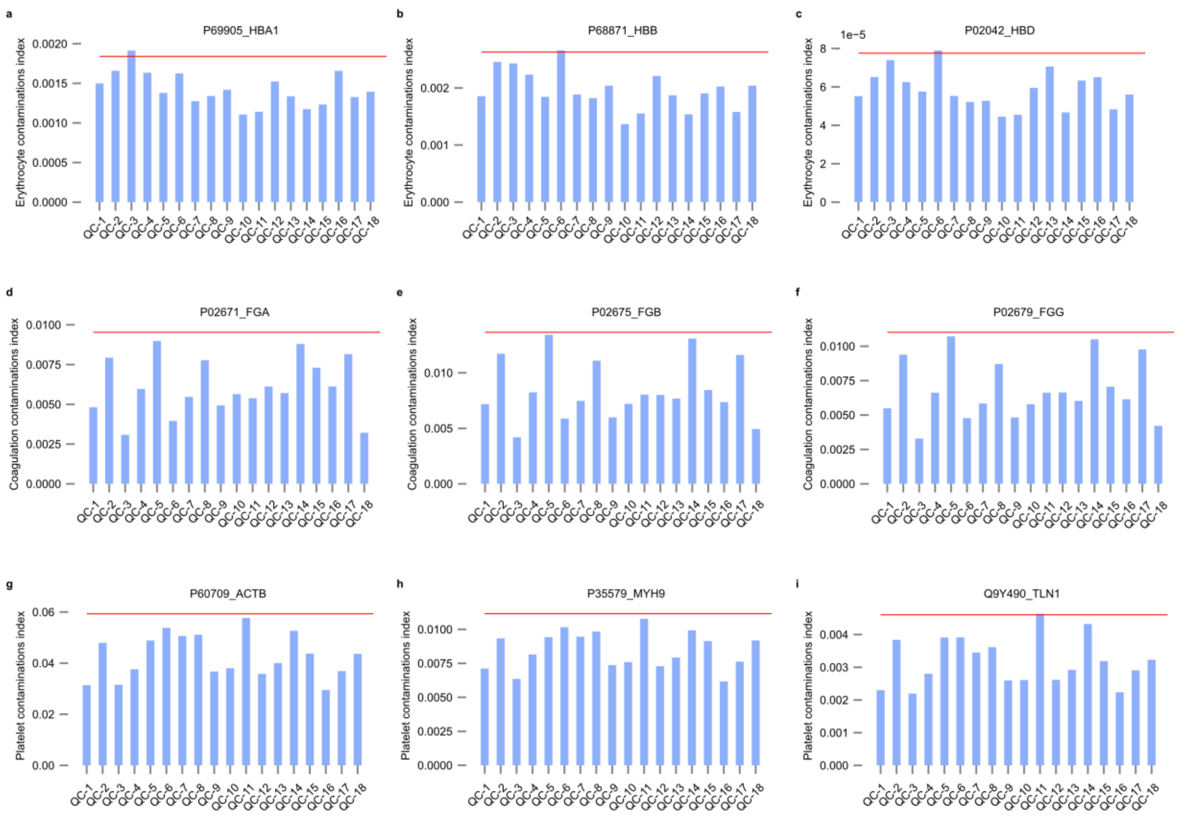

**Supplemental Figure. 5 Assessment of potential contamination of 18 QC samples in Project 1.** Samples with values exceeding ±2 standard deviations from the mean (horizontal red lines) are flagged as potentially contaminated.

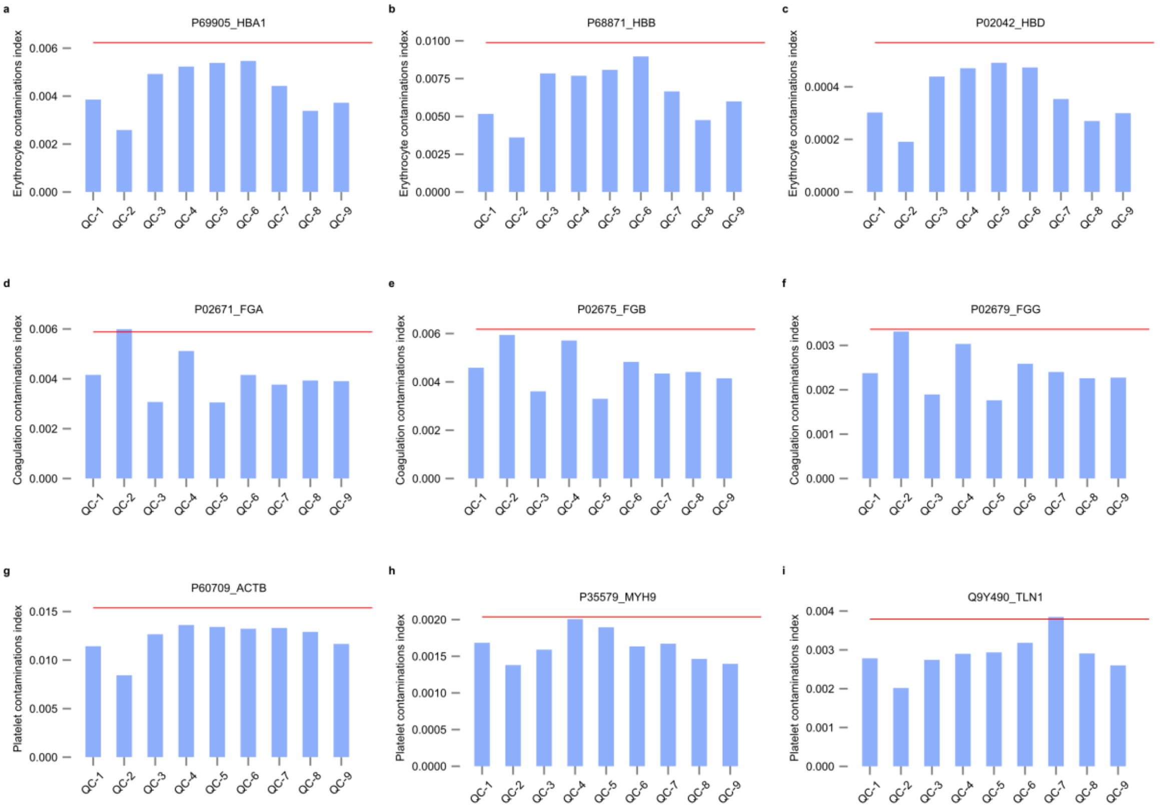

**Supplemental Figure. 6 Assessment of potential contamination of 9 QC samples in Project 2.** The horizontal red lines represent the mean ±2 standard deviations, which are used as a threshold to identify potentially contaminated samples.

### 7. Proteonano preserves quantitative accuracy

The quantitative accuracy of the Proteonano workflow was assessed by adapting an established assay[2,3]. Pooled healthy donor plasma (Ori Biotech, Shanghai, China) was generated by mixing plasma from whole blood samples collected in K2-EDTA containing tubes. *Saccharomyces cerevisiae* (*S. cerevisiae*) was lysed in 100 mM HEPES pH 7.4, 150 mM KCl, and 1 mM $MgCl_2$ by passing through a gauge 12 syringe 15 times on ice, followed by filtration (0.2 µm). *Escherichia coli* (*E. coli*) was homogenized and lysed, then filtered (0.2 µm). Protein concentration for each sample was determined using a UV spectrometer at 205 nm (Nano 300, Allsheng Instruments, China). Each sample was then mixed with fixed ratios of *E. coli* and *S. cerevisiae*, resulting in 2:1 and 1:2 fold changes, respectively. For conditions A and B, 40 µL of plasma

(~1000 µg proteins) was spiked with 2.5 or 5 µg of *S. cerevisiae* and 5 or 2.5 µg of *E. coli* lysate, respectively. Samples with the reference neat digestion workflow or the Proteonano workflow were subjected to preparation followed by LC-MS/MS measurements.

In this experiment, Sample A contained twice the amount of *E. coli* proteins compared to Sample B, while Sample B had twice the amount of *S. cerevisiae* proteins relative to Sample A. The amount of pooled healthy donor plasma proteins remained constant across all samples. We conducted proteomic data analysis according to a previous study[3]. The log-transformed ratios (Log2(A/B), y-axis) of protein intensities were plotted against the log-transformed intensities of proteins in Sample B (Log2(B), x-axis), as presented in Supplemental Fig. 7. The theoretical values for Log2(A/B) were fixed at 1, 0, and -1 for proteins derived from *E. coli*, human plasma, and *S. cerevisiae*, respectively. The results show that the colored circles representing the proteins were distributed around their respective theoretical reference lines (black horizontal dashed lines in Fig. 7). Moreover, linear regression analysis revealed that the proteins detected using the Proteonano workflow were closer to the reference lines across all three species, indicating improved quantification accuracy.

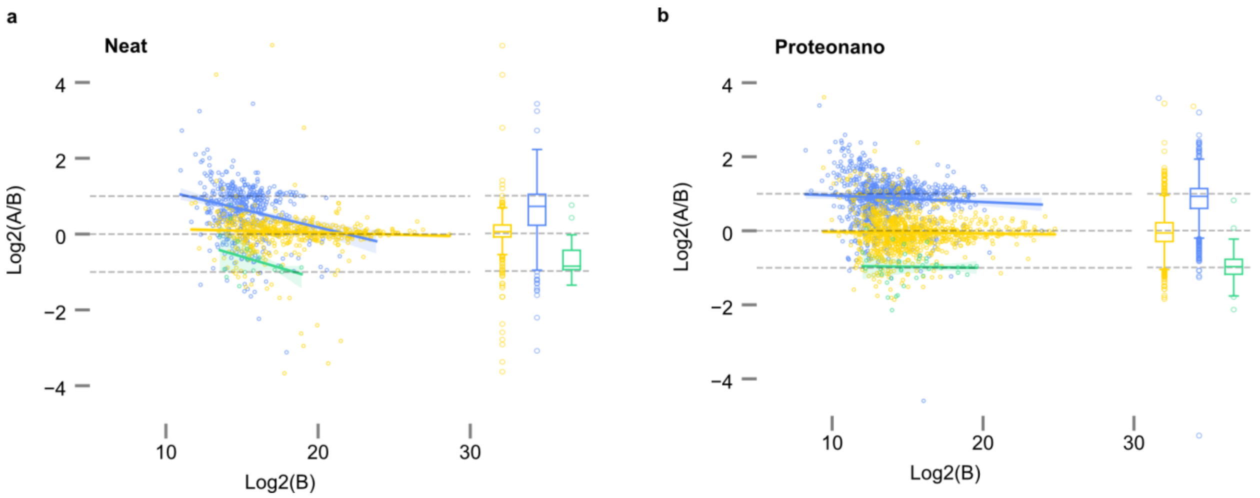

**Supplemental Figure. 7:** Comparison of relative protein abundance detected using the Proteonano workflow and neat plasma workflow. Log-transformed ratios (log2(A/B), y-axis) of proteins from three species are plotted against the log-transformed intensity of proteins in sample B (Log2(B), x-axis). Black horizontal dashed lines indicate the expected log2(A/B) values for proteins from human (yellow circles), *S. cerevisiae* (green circles), and *E. coli* (blue circles). Solid lines represent the local trends along the x-axis for the experimental log-transformed ratios of each population, with yellow, green, and blue lines corresponding to human plasma, *S. cerevisiae,* and *E. coli,* respectively. **a**. The neat plasma digestion workflow. **b**. The Proteonano workflow.

### 8. Fold-change accuracy of the Proteonano workflow

The experiments were conducted according to previous research[4]. To be brief, total protein concentrations of human and porcine plasma were determined by the Micro BCA Protein Assay Kit (ThermoFisher Scientific). Human and porcine plasma were mixed at different proportions at a final concentration of 14 mg/ml, corresponding to total protein concentration in 1:5 diluted human plasma. 100 uL of mixed human/porcine plasma samples with 11 distinct ratios (1:0, 99:1, 9:1, 2:1, 1.5:1, 1:1, 1:1.5, 1:2, 1:9, 1:99, 1:0, w/w) were processed either by the Proteonano workflow or the reference neat digestion workflow. For LC-MS/MS measurement, 300 ng of peptides of each sample were separated by Vanquish™ Neo UHPLC system (ThermoFisher Scientific) coupled to an Orbitrap Astral mass spectrometer (ThermoFisher) with a throughput of 100SPD. Spectra were searched using DIA-NN software (version 1.8.1) in library free mode against a protein library containing UniProt *homo sapiens* reviewed entries (20,422 entries) and *Sus scrofa* reviewed entries (3,590 entries). Protein intensities were then used to calculate ratios of detected proteins in samples at different mixing proportions, i.e. measured fold-changes.

The fold-change accuracy of Proteonano workflow was assessed by the spike-in experiment. Pearson correlation was calculated to compare the observed and expected fold-changes of human proteins in mixed plasma samples. At a correlation threshold exceeding 0.8, the Proteonano workflow identified 148 human protein groups, while the neat digestion workflow (Neat) detected only 19 protein groups (Supplementary Fig. 8b). We then evaluated the fold-change accuracy by comparing the expected and observed fold-changes in human/porcine mixed plasma samples. Supplementary Fig. 8c illustrates this comparison, where yellow dots (left) and blue dots (right) represent the observed fold-changes (x-axis) for protein groups identified using the neat workflow and Proteonano, respectively. Only protein groups with a Pearson correlation coefficient exceeding 0.80 in the neat workflow or 0.90 in the Proteonano workflow are included. The observed fold-changes for each protein are connected by light yellow (Neat) and blue dashed lines (Proteonano). Using linear regression to fit the data (observed fold-changes versus expected fold-changes), we obtained slopes of 0.701 and 0.762 for the Neat and Proteonano workflows, respectively. Notably, a slope closer to 1 indicates better fold-change accuracy.

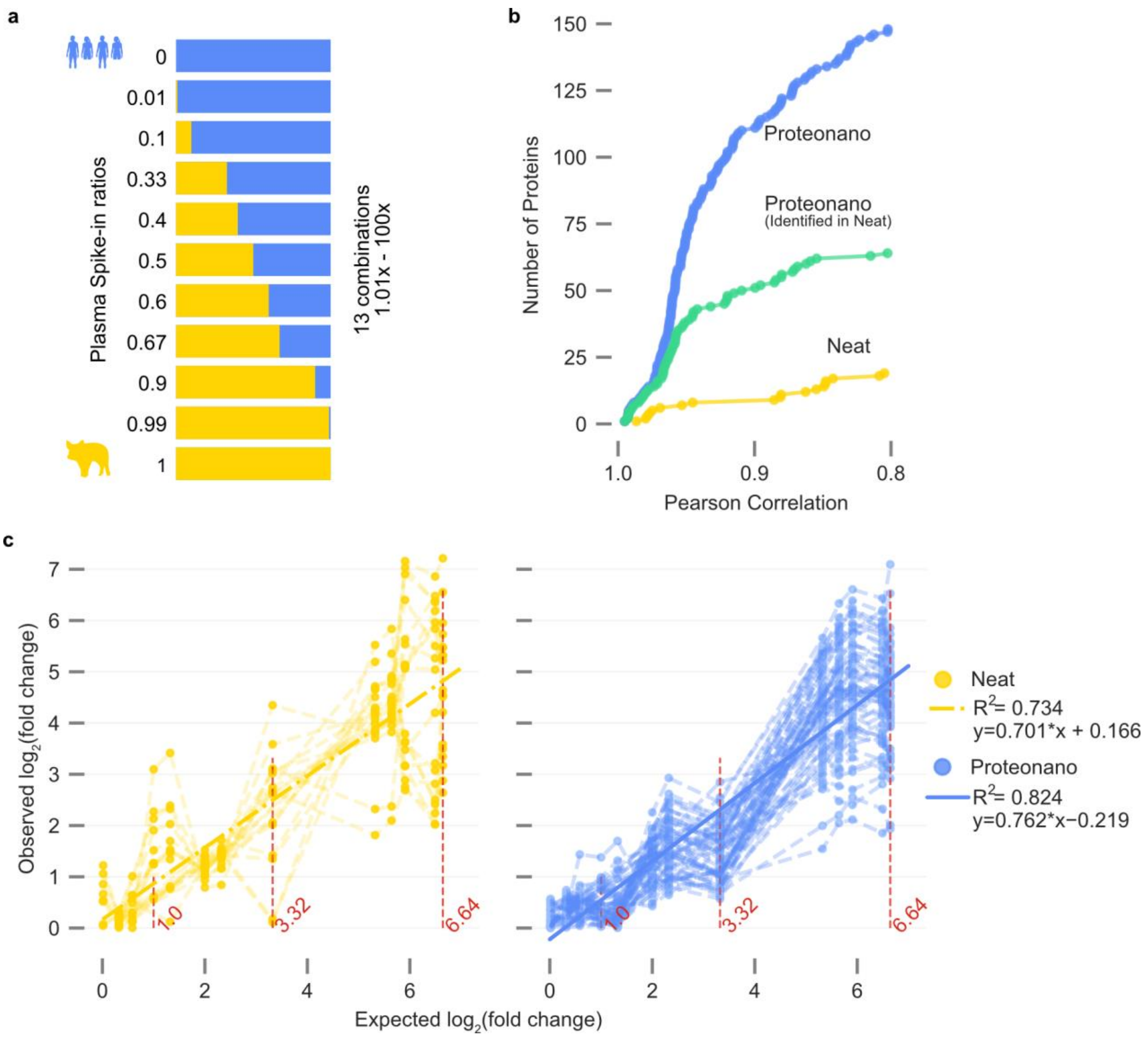

**Supplemental Figure. 8 Fold-change accuracy of protein quantification for neat and the Proteonano workflow. a** Schematic of the spike-in experiment, where a bovine plasma proteome is introduced into a human plasma proteome at seven distinct ratios. **b** Human proteins identified at varying Pearson correlation thresholds are compared. The X-axis represents Pearson correlation values (truncated at 0.8), while the Y-axis displays the number of human proteins exceeding each threshold. Shared protein groups detected in both workflows are marked green. Proteonano identifies 184 human protein groups (blue dots, Pearson correlation > 0.8) compared to 19 identified by the neat workflow (yellow dots, Pearson correlation > 0.8). **c** Fold-change accuracy between the expected log2(fold-change) (x-axis) and observed log2(fold-change) (y-axis) between human/porcine mixed plasma samples. The red vertical dash lines denote the expected log2(fold-changes) at 1.0, 3.32, 6.64. Yellow dots (left) and blue dots (right) represent the observed fold changes of the protein groups identified by the neat and Proteonano, respectively. Only protein groups exhibiting a Pearson correlation coefficient greater than 0.80 detected in the neat workflow or greater than 0.90 for the Proteonano workflow are shown. Light yellow and blue dashed lines connect the observed fold-changes of each protein. Dark yellow dashed line (left) and dark blue solid line (right) represent the linear regression between the observed log2(fold-changes) (y-axis) and the expected log2(fold-changes) (x-axis) of the protein groups included in the Neat and Proteonano workflow, respectively. The slope of the linear regression is estimated to be 0.701 (Neat) or 0.762 (Proteonano).

### 9. Supplementary information on the proof-of-concept study

**Supplemental Table 3.** Patient characteristics of the showcase cohort.

| | **Overall** | **Cognitive Decline** | **Control** | **P-Value** |
|---|---|---|---|---|
| **N** | 183 | 58 | 125 | |
| **Age, median [Q1, Q3]** | 72.0 [69.0,77.0] | 75.0 [68.0,81.0] | 72.0 [69.0,75.0] | 0.061 |
| **Gender, n (male %)** | 68 (37.2) | 20 (34.5) | 48 (38.4) | |

**Supplemental Table 4.** The list of the identified low abundance DEPs.

| **Uniport ID** | **Protein names** | **Gene Names** | **HPPP (ng/mL)** |
|---|---|---|---|
| P07203 | Glutathione peroxidase 1 | GPX1 | 3.8 |
| P14770 | Platelet glycoprotein IX | GP9 | 4.3 |
| P15153 | Ras-related C3 botulinum toxin substrate 2 | RAC2 | 4.2 |
| P20160 | Azurocidin | AZU1 | 6.6 |
| P26447 | Protein S100-A4 | S100A4 | 9.3 |
| P53597 | Succinate--CoA ligase subunit alpha, mitochondrial | SUCLG1 | 0.17 |
| P59998 | Actin-related protein 2/3 complex subunit 4 | ARPC4 | 6.4 |
| P60953 | Cell division control protein 42 homolog | CDC42 | 2.8 |

| | | | |
|---|---|---|---|
| P63000 | Ras-related C3 botulinum toxin substrate 1 | RAC1 | 3.6 |
| Q01082 | Spectrin beta chain, non-erythrocytic 1 | SPTBN1 | 2.4 |
| Q15365 | Poly(rC)-binding protein 1 | PCBP1 | 2.3 |
| Q6PK18 | 2-oxoglutarate and iron-dependent oxygenase domain-containing protein 3 | OGFOD3 | 0.015 |
| Q8N392 | Rho GTPase-activating protein 18 | ARHGAP18 | 2.1 |

### 10. Effect of proteomic search reference libraries

The size of reference libraries can influence the number of protein groups identified in MS-based proteomic analyses. To evaluate this effect, we subjected MS data from Orbitrap Astral mass spectrometer, following Proteonano workflow, to searches against three reference libraries of varying sizes: Swiss-Prot (~20,000 proteoforms), Proteomes (~80,000 proteoforms), and TrEMBL (~200,000 proteoforms). The searches yielded only modest increases in identified protein groups within the same MS dataset (Supplementary Fig. 9), with the TrEMBL library identifying ~15% more protein groups compared to the Swiss-Prot library. However, given that Swiss-Prot includes only validated proteoforms while TrEMBL encompasses a substantial number of unvalidated proteoforms, the benefits of using the larger TrEMBL library remain limited.

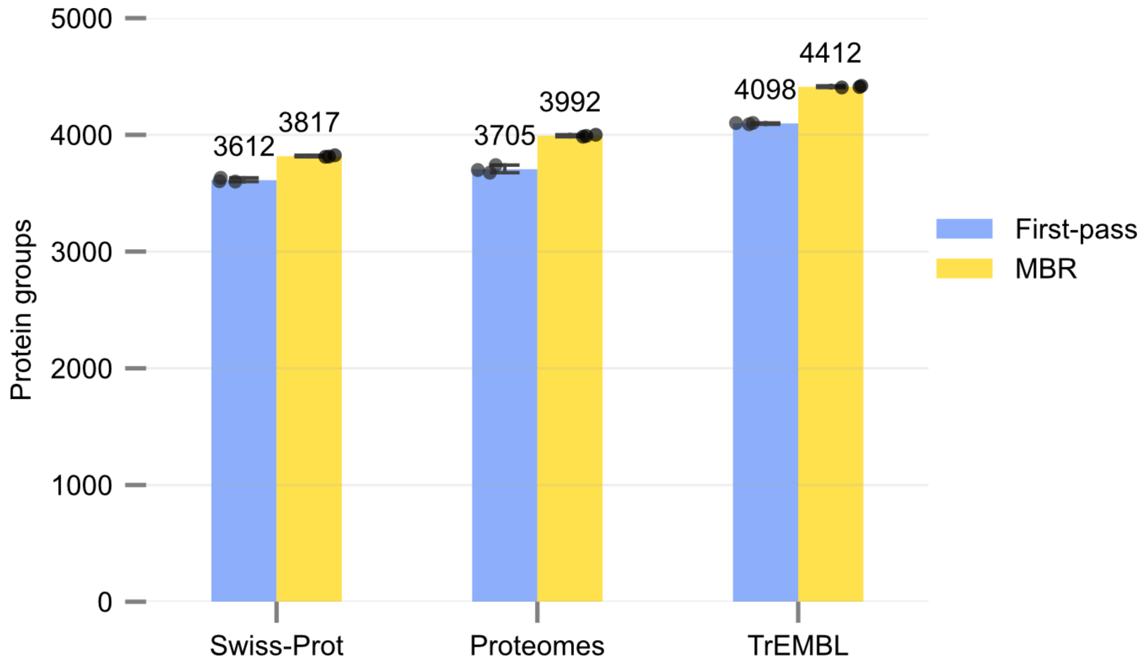

**Supplemental Figure. 9:** Effect of reference databases on identified protein groups.

### 11. Assessment of sample related factors

It is known that plasma hemolysis could impact numbers of protein groups detected by MS-based proteomics[5,6]. However, if this could affect proteomic detection in samples processed by Proteonano remains unknown. To test this, we first determined the effect of hemolytic state of plasma samples. Plasma samples with different extents of hemolysis (graded as 0, no hemolysis (n=3), 1, some hemolysis (n=5), 2, extensive hemolysis (n=2), based on color of plasma samples) were centrifuged, and processed by the Proteonano workflow. Analysis using a timsTOF Pro2 mass spectrometer revealed that the number of protein groups identified increased with the severity of hemolysis in the sample (Supplementary Fig. 10). This trend is consistent with findings from samples processed using conventional methods, suggesting that blood cell protein contamination similarly affects the Proteonano workflow for plasma proteomic sample preparation.

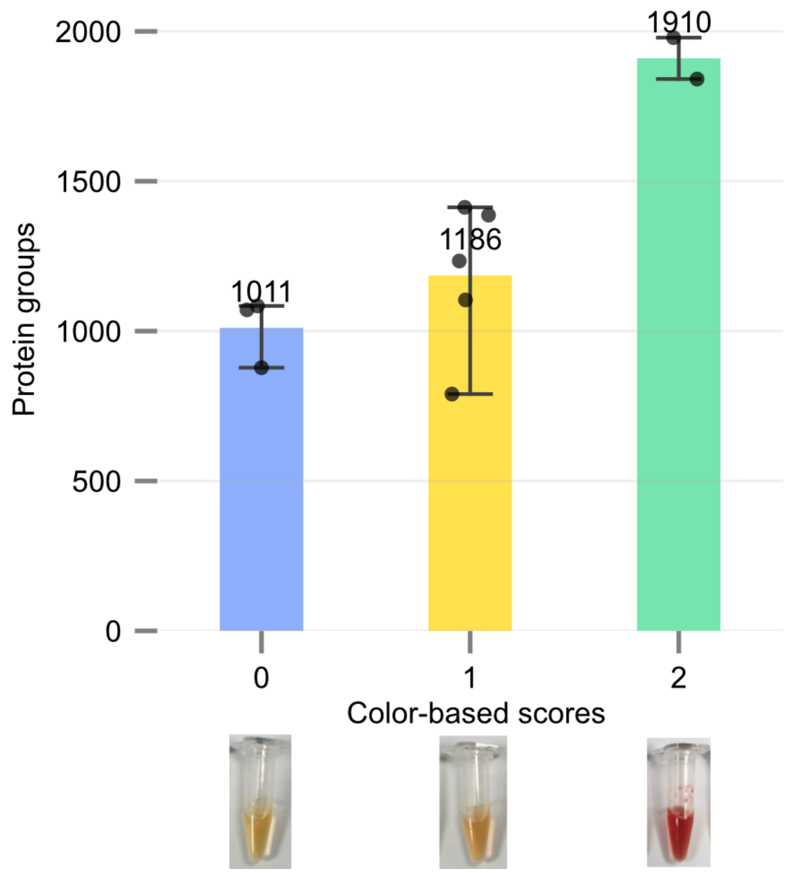

**Supplemental Figure. 10:** Detection depth of samples with various levels of hemolysis. Samples were processed by the Proteonano platform and analyzed by a timsTOF Pro2 mass spectrometer.

### 12. Parameters on the LC-MS/MS methods

#### Liquid chromatography setups

LC columns used include: PepMap™ Neo Trap Column, 5 µm C18 300 µm x 5 mm (ThermoFisher Scientific 174500) EASY-Spray™ Column, 2 µm C18 150 µm x 15 cm (ThermoFisher Scientific ES906) EASY-Spray™ PepMap™ Neo Column, 2 µm C18 75 µm x 50 cm (ThermoFisher Scientific ES75500) µPAC™ Neo Column, 2.5 µm x 16 µm, 110 cm (ThermoFisher Scientific COL-NANO110NEOB).

After HPLC separation, samples were fed into an Orbitrap Astral Mass Spectrometer (ThermoFisher Scientific) with a fused silica spray needle (ThermoFisher Scientific EV1111) and EasySpray adapter (ThermoFisher Scientific EV-1072).

**Supplemental Table 5.** The SPD method.

| 180 SPD method (Trap/Elute), ES906 chromatography column | | | |
|---|---|---|---|
| Time/min | Duration/min | %B | Flow rate / $mL \cdot min^{-1}$ |
| 0.0 | 0.0 | 4.0 | 2.5 |
| 4.0 | 4.0 | 25.0 | 2.5 |
| 5.8 | 1.8 | 35.0 | 2.5 |
| Column wash | | | |
| 6.2 | 2.5 | 99.0 | 2.5 |
| 6.9 | 2.5 | 99.0 | 2.5 |
| Stop run | | | |
| Column equilibration | | | |

**Supplemental Table 6.** The 100SPD method.

| 100 SPD method (Trap/Elute), ES906 chromatography column | | | |
|---|---|---|---|
| Time/min | Duration/min | %B | Flow rate / $mL \cdot min^{-1}$ |
| 0.0 | 0.0 | 1.0 | 1.8 |
| 0.7 | 0.7 | 4.0 | 1.8 |
| 1.0 | 0.3 | 8.0 | 1.8 |
| 7.7 | 6.7 | 25.0 | 1.8 |
| 11.4 | 3.7 | 35.0 | 1.8 |
| 11.8 | 0.4 | 55.0 | 2.5 |
| Column wash | | | |
| 12.3 | 0.5 | 99.0 | 2.5 |
| 13.0 | 0.7 | 99.0 | 2.5 |
| Stop run | | | |

| |
|---|
| Column equilibration |

**Supplemental Table 7.** The 60SPD method.

| 60 SPD method (Trap/Elute), ES906 chromatography column | | | |
|---|---|---|---|
| Time/min | Duration/min | %B | Flow rate / mL·min$^{-1}$ |
| 0.0 | 0.0 | 4.0 | 2.0 |
| 0.5 | 0.5 | 5.0 | 2.0 |
| 0.9 | 0.4 | 8.5 | 0.8 |
| 13.9 | 13.0 | 25.0 | 0.8 |
| 20.8 | 6.9 | 35.0 | 0.8 |
| 21.2 | 0.4 | 55.0 | 2.0 |
| Column wash | | | |
| 21.7 | 0.5 | 99.0 | 2.0 |
| 22.6 | 0.9 | 99.0 | 2.0 |
| Stop run | | | |
| Column equilibration | | | |

**Supplemental Table 8.** The 24SPD method

| 24 SPD method (Trap/Elute), ES906 chromatography column | | | |
|---|---|---|---|
| Time/min | Duration/min | %B | Flow rate / mL·min$^{-1}$ |
| 0.0 | 0.0 | 4.0 | 2.5 |
| 0.5 | 0.5 | 5.0 | 2.5 |
| 1.0 | 0.5 | 7.0 | 0.6 |
| 39.1 | 38.1 | 20.0 | 0.6 |
| 57.1 | 18.0 | 35.0 | 0.6 |

| | | | |
|---|---|---|---|
| 57.4 | 0.3 | 55.0 | 2.5 |
| Column wash | | | |
| 57.9 | 0.5 | 99.0 | 2.5 |
| 58.6 | 0.7 | 99.0 | 2.5 |
| Stop run | | | |
| Column equilibration | | | |

**Supplemental Table 9.** MS parameters utilized in all experiments

| | **Property** | **Setting** |
|---|---|---|
| **Method setting** | Application mode | Peptide |
| **Ion source** | Positive Ion (V) | 2000 |
| | Ion Transfer Tube Temp (°C) | 275 |
| **MS global setting** | Advanced Peak Determination | TRUE |
| | Default Charge State | 2 |
| **Orbitrap analyzer full scan** | Scan Range (m/z) | 380-980 |
| | Detector Type | Orbitrap |
| | Orbitrap Resolution | 240000 |
| | Max IT (ms) | 5 |
| | RF Lens (%) | 40 |
| | AGC Target (%) | 500 |
| **Astral analyzer DIA MS2 scan** | Scan Range (m/z) | 150-2000 |
| | Isolation Window (m/z) | 2 |
| | Windows Overlap (m/z) | 0 |
| | Window Placement Optimization | On |
| | Number of Scan Events | 300 |

| | | |
|---|---|---|
| | HCD Collision Energies (%) | 25 |
| | Detector Type | Astral |
| | Max IT (ms) | Experiment Dependent |
| | AGC Target (%) | 500 |
| | Loop Control | Time |
| | Loop Time (sec) | 0.6 |

**Supplemental Table 10.** Maximum ion injection time for each SPD method

| **SPD method Maximum injection time** | |
|---|---|
| 180 SPD | 3.0 ms |
| 100 SPD | 3.5 ms |
| 60 SPD | 5.0 ms |
| 24 SPD | 7.0 ms |
| 15 SPD | 7.0 ms |
| 14 SPD | 7.0 ms |
| 11 SPD | 7.0 ms |
| 7 SPD | 7.0 ms |

**Supplemental Table 11. The** 14SPD method.

| 14S PD method (Direct Infusion), ES75500 column | | | |
|---|---|---|---|
| Time/min | Duration/min | %B | Flow rate / mL·min$^{-1}$ |
| 0.0 | 0.0 | 5.0 | 0.3 |
| 55.0 | 55.0 | 25.0 | 0.3 |
| 65.0 | 10.0 | 35.0 | 0.3 |

| | | | |
|---|---|---|---|
| Column wash | | | |
| 70.0 | 5.0 | 99.0 | 0.3 |
| 80.0 | 10.0 | 99.0 | 0.3 |
| Stop run | | | |
| Column equilibration | | | |

**Supplemental Table 12.** The 15SPD method with µPac110cm column.

| 15 SPD method (Direct Infusion), µPac110cm column | | | |
|---|---|---|---|
| Time/min | Duration/min | %B | Flow rate / mL·min$^{-1}$ |
| 0.00 | 0.00 | 4.00 | 0.75 |
| 0.40 | 0.40 | 4.00 | 0.75 |
| 47.40 | 47.00 | 22.50 | 0.75 |
| 60.40 | 13.00 | 45.00 | 0.75 |
| Column wash | | | |
| 64.90 | 4.50 | 99.00 | 0.75 |
| 66.50 | 1.60 | 99.00 | 0.75 |
| Stop run | | | |
| Column equilibration | | | |

**Supplemental Table 13.** The 11SPD method with µPac110cm column.

| 11 SPD method (Direct Infusion), µPac110cm column | | | |
|---|---|---|---|
| Time/min | Duration/min | %B | Flow rate / mL·min$^{-1}$ |
| 0.00 | 0.00 | 1.00 | 0.40 |
| 0.10 | 0.10 | 2.00 | 0.40 |
| 75.10 | 75.00 | 22.50 | 0.40 |

| | | | |
|---|---|---|---|
| 92.10 | 17.00 | 45.00 | 0.40 |
| Column wash | | | |
| 97.60 | 0.75 | 99.00 | 0.75 |
| 100.00 | 0.75 | 99.00 | 0.75 |
| Stop run | | | |
| Column equilibration | | | |

**Supplemental Table 14.** The 7SPD method with µPac110cm column.

| 7 SPD method (Direct Infusion), µPac110cm column | | | |
|---|---|---|---|
| Time/min | Duration/min | %B | Flow rate / mL·min$^{-1}$ |
| 0.00 | 0.00 | 1.00 | 0.25 |
| 0.10 | 0.10 | 2.00 | 0.25 |
| 110.10 | 110.00 | 22.50 | 0.25 |
| 150.10 | 40.00 | 45.00 | 0.25 |
| Column wash | | | |
| 150.60 | 0.50 | 99.00 | 0.25 |
| 175.00 | 24.40 | 99.00 | 0.25 |
| Stop run | | | |
| Column equilibration | | | |
